# Lipidomic analysis of adipose-derived extracellular vesicles reveals their potential as lipid mediators of obesity-associated metabolic complications

**DOI:** 10.1101/2021.12.10.472057

**Authors:** Alexia Blandin, Grégory Hilairet, Maharajah Ponnaiah, Simon Ducheix, Isabelle Dugail, Bertrand Cariou, Marie Lhomme, Soazig Le Lay

## Abstract

Adipose extracellular vesicles (AdEV) transport lipids that could participate to the development of obesity-related metabolic dysfunctions. This study aimed to define mice AdEV lipid signature in either healthy or obesity context by a targeted LC-MS/MS approach.

Distinct clustering of AdEV and visceral adipose tissue (VAT) lipidomes by principal component analysis reveals specific lipid composition of AdEV compared to source VAT. Comprehensive analysis identifies enrichment of ceramides and phosphatidylglycerols in AdEV compared to VAT in lean conditions. Lipid subspecies commonly enriched in AdEV highlight specific AdEV-lipid sorting. Obesity impacts AdEV lipidome, driving triacylglycerols and sphingomyelins enrichment in obese versus lean conditions. Obese mice AdEV also display elevated phosphatidylglycerols and acid arachidonic subspecies contents highlighting novel biomarkers and/or mediators of metabolic dysfunctions.

Our study identifies specific lipid-fingerprints for plasma, VAT and AdEV that are informative of the metabolic status and underline the signaling capacity of lipids transported by AdEV in obesity-associated complications.

## Introduction

Epidemic obesity, with nearly 40 % of the world’s adult population being overweight or obese, is the greatest threat to global health, by increasing risk of type 2 diabetes (T2D), cardiovascular and liver diseases. Multiple factors (environmental, genetic and biological) interact to cause obesity. Adipose tissue (AT) hypertrophy and consecutive metabolic dysfunction specifically lead to systemic lipid overflow, lipotoxic fat depots in peripheral organs and low-grade inflammation via dysregulated production of adipokines, which, altogether, participate in the settings of metabolic complications. Recent evidences indicate that a significant part of the AT secretome is in the form of AT-derived extracellular vesicles (AdEV), that may contribute to the development of obesity-related metabolic complications (1, 2).

Others and we previously demonstrated significant increase of plasma EV concentrations in patients with obesity, with a strong positive association with HOMA-IR indicating insulin-resistance (IR) and subsequent T2D risk (3, 4). AdEV are viewed as critical mediators of metabolic alterations since the injection of AdEV derived from obese AT into healthy mice triggers IR (5, 6). Efforts to understand how AdEV promote metabolic dysfunction have focused on their protein or miRNA composition and subsequent transfer to recipient cells (5–7). However, little attention was paid to their potential as lipid species carriers even if some lipids, like ceramides or diacylglycerols (DAG), are recognized as potent mediators of IR development in skeletal muscle or liver (8, 9).

High-resolution lipidomic analysis applied on EV from different cell sources identified up to 2,000 different lipid species (10, 11), which provided a basis for structural properties which likely participate in EV stability in biofluids (12). A recent study also pointed out that adipocytes can release neutral lipid-filled EV, via a lipase-independent pathway distinct from that used in canonical lipid mobilization by lipolytic release of free fatty acids (13). Others highlight the participation of AdEV in the transport of free fatty acids fueling melanoma aggressiveness with energetic substrates (14). In line, studies focusing on non-adipose derived EV demonstrated that palmitate-induced EV were enriched in ceramides, supporting the idea that EV contribute to sphingolipid efflux pathway (15, 16). Moreover, EV can transfer ceramides to macrophages or muscle cells thereby inducing IR in recipient cells (17, 18). Previous lipidomic studies on cultured 3T3-L1 adipocyte-derived EV identified a predominance of phospholipids, sphingolipids and traces of glycerolipids, reflecting the composition of adipocyte plasma membrane (19, 20). We could also appreciate subtle differences in EV lipid fingerprint depending on EV subtype, namely large EV (lEV) shed from the plasma membrane and small EV (sEV) which originate from the endosomal system (10, 11, 21). Our previous work highlighted a specific cholesterol enrichment in 3T3-L1 adipocyte sEV in agreement with the role of this sterol in EV biogenesis, whereas adipocyte IEV presented high amounts of externalized phosphatidylserine (PS) in line with the pro-coagulant potential of this EV subclass (19).

Considering the role assigned to AdEV as lipid transporters and as mediators of obesity-related metabolic complications, we aimed to compare AdEV lipid content, with respect to secreting AT and circulating lipids, in the lean and pathophysiological context of obesity. To this purpose, we performed a targeted lipidomic analysis to compare the lipidome of sEV and IEV with source visceral AT (VAT) and with plasma in lean and obese (*ob/ob*) mice. We present here comprehensive lipid maps revealing specific adipose EV lipid sorting when compared to secreting VAT. We demonstrated that AdEV lipidome is more dependent on VAT pathophysiological state rather than on EV subtype and we identified some specific AdEV lipid classes or species closely related to the obese state. Particularly, enrichment in some EV lipid subspecies may constitute novel candidates/mediators in metabolic dysfunctions associated with obesity.

## Methods

### Animal experimentation

Adult mice heterozygous (*Ob*/+) for the leptin spontaneous mutation Lep^ob^ were initially obtained from Charles River (JAX™ mice strain) and interbred to obtain a colony. Regular backcross with commercial Lep^ob/+^ is performed to avoid any background drift. At 3-month of age, *ob/ob* animals were identified on the basis of their increased body weight that associates with hyperglycemia, hyperinsulinemia and significant increase of liver and AT mass at the expense of muscle mass (Table S1).

Three-month old lean or obese mice were used to collect VAT explants for EV isolation. We retained only male mice since sex-specific lipid signature has been described in *ob/ob* mice (22). All mice had *ad libitum* access to food and water and were housed in the same open mouse facility on a day/night cycle. Animals were killed in a non-fasted state.

Animal care and study protocols were approved by the French Ministry of Education and Research and the ethics committee N°6 in animal experimentation and were in accordance with the EU Directive 2010/63/EU for animal experiments.

### Plasma collection

Mice peripheral blood was collected on EDTA-coated tubes following intracardiac puncture. Platelet-rich plasma was separated from whole blood by a 5 min centrifugation at 1,500 x*g*, and recentrifuged for 5 min at 1,900 x*g* to obtain platelet-free plasma (PFP).

### VAT-derived EV isolation

Mice VAT were minced into small pieces (50-150 mm^3^) and were placed into Clinicell^®^ 25 cassettes (Mabio, France) filled with 10mL ECBM/Hepes 10mM/BSA FFA free 0,1% pH 7.4 as previously described (23). Serum-free conditioned medium (CM) after 48h culture was collected, filtered on 100μm cell strainers and use for EV isolation similarly to our previous characterization of adipocyte-derived EV reported on EV Track knowledgebase (24) (http://evtrack.org/, ID: EV210202). The absence of serum prevented AdEV preparations from contamination by external bovine source of EV. IEV were recovered from cell-cleared supernatants (1,500 x*g* for 20 min) by centrifugation 1 hour at 13,000 x*g*, followed by two washing steps in NaCl and resuspended in sterile NaCl. sEV were further isolated from lEV-depleted supernatants following a 100,000 x*g* ultracentrifugation step for 1 hour at 4°C (rotor MLA-50, Beckman Coulter Optima MAX-XP Ultracentrifuge) and two washes in NaCl before resuspension in NaCl.

### Nanoparticle Tracking Analysis

EV samples were diluted in sterile NaCl before nanoparticle tracking analysis (NTA). NTA was undertaken using the NanoSight NS300 (Malvern Instruments, Malvern, UK) equipped with a 405 nm laser. Ninety-second videos were recorded in five replicates per sample with optimized set parameters (the detection threshold was set to 5 for both EV subtypes). Temperature was automatically monitored and ranged from 20°C to 21°C. Videos were analyzed when a sufficient number of valid trajectories was measured. Data capture and further analysis were performed using the NTA software version 3.1. EV concentrations are expressed as number of particles secreted by adipocytes, the number of adipocytes present in the secreting VAT being estimated by indirect calculation as we previously described (19).

### Western Blotting

VAT explants were resuspended in lysis buffer [50 mM Tris pH 7.4, 0.27 M sucrose, 1 mM Na-orthovanadate pH 10, 1 mM ethylenediaminetetraacetic acid (EDTA), 1 mM ethylene glycol-bis(β-aminoethyl ether)-N,N,N’,N’-tetraacetic acid (EGTA), 10mMNa β-glycerophosphate, 50mM NaF, 5 mM Na pyrophosphate, 1% (w/v) Triton X-100, 0.1% (v/v) 2-mercaptoethanol and cOmplete™ Protease Inhibitor Cocktail (Roche Diagnostics). Whole cell lysates were centrifuged at 13 000 x*g* for 10min at 4°C to get rid of insoluble material. Isolated AdEV following differential centrifugation were resuspended in NaCl. AdEV and VAT protein content was estimated by DC-protein assay (BioRad) by using BSA as standard. Protein lysates were stored at −20°C until use.

8μg of protein lysates were diluted in Laemli Buffer 4X (Biorad) in reducing conditions, heated at 95°C for 10 min and migrated on a 4–15% Mini-Protean TGX gel (Biorad) and transferred on to nitrocellulose membranes using Trans Blot Turbo apparatus (Biorad). Membranes were blocked for 90 min at room temperature using TBS blocking buffer (LI-COR Biosciences) and incubated with primary antibodies diluted in the same blocking buffer. Antibodies used for Western-blot were previously detailed (19). IRDye secondary antibodies (LI-COR Biosciences) were used for protein detection and digital fluorescence was visualized by an Odyssey CLX system (LI-COR Biosciences). Immunoblot quantification was performed following analysis of protein signal by Image Studio^®^ software.

### Transmission Electron Microscopy (TEM)

IEV and sEV were fixed for 16 h at 4°C with 2.5% glutaraldehyde (LFG Distribution, Lyon, France) in 0.1 M Sorensen buffer pH 7.4 then deposited on formwar-coated copper grids and negatively stained with phosphotungstic acid 1% (w/v) for 30 seconds. Grids were rinsed with milliQ water, let to air dry and observed with a Jeol JEM 1400 microscope (Jeol, France) operated at 120 KeV.

### Lipidomic analyses

All lipidomics analyses were performed on the ICANalytics platform (IHU ICAN, Paris, France). EV lipidomics data originated from 2 independent batches (n=10 lean and n=8 obese), whereas tissue and plasma lipidomics data originated from one batch only (n=4-5 obese, n=4-5 lean). In order to adjust for batch effect in EV, data were combined using a relative difference method with Multi Experiment Viewer (MeV) software version 4.9 (https://sourceforge.net/projects/mev-tm4/) (25). Delta was calculated between the comparison group, and the subsequent relative values were processed for the statistical analysis.

#### Tissue homogenization

VAT were weighted and supplemented with isopropanol to a final concentration of 80mg/mL. Tissues were homogenized using ceramic beads and the “soft” program of the Precellys Evolution instrument (Bertin Instruments, France).

#### Lipid extraction

Lipids were extracted from 10μl platelet-free plasma, 50μl EV or 4mg VAT lysate using a modified Bligh and Dyer method. Samples were supplemented with deuterated or odd chain internal standards (CE(18:1d7), cholesterol d7, cer(d18 :1/24 :0d7), LPC(17:1), LPE(18:1d7), PA(15:0/18:1d7), PC(16:0/16:0d9), PC(15:0/18:1d7), PE(15:0/18:1d7), PG(15:0/18:1d7), PI(17:0/20:4), PS(16:0/18:1d31), SM(d18:1/16:0d31), TAG(17:0/17:1/17:0 d5), DG(15:0/18:1d7) from Avanti Polar Lipids) serving for quantification of the endogenous lipid species. Lipids were extracted with 1.2mL methanol/CHCl3 (2:1 v/v) in the presence of the antioxidant BHT and 310μl HCl 0.005N. Phase separation was triggered by addition of 400μl CHCl_3_ and 400μl water, followed by a centrifugation 3,600 *xg* for 10min at 4°C. The organic phase was harvested, was dried and then resuspended in 40μl of LC/MS solvent (Chloroform/acetonitrile/Isopropanol (80:19:1 v/v/v)). A control plasma was extracted in parallel and injected every 10 samples to correct for signal drift.

#### LC-MS/MS analysis of phospholipids and sphingolipids

Lipids were quantified by LC-ESI/MS/MS using a prominence UFLC and a QTrap 4000 mass spectrometer. Sample (4μl) was injected to a kinetex HILIC 2.6μm 2.1×150mm column. Mobile phases consisted of water and acetonitrile containing 30mM ammonium acetate and 0.2% acetic acid. Lipid species were detected using scheduled multiple reaction monitoring (sMRM) in the positive-ion mode reflecting the headgroup fragmentation of each lipid class.

#### LC-MS/MS analysis of neutral lipids (CE, FC, DAG and TAG)

Sample (4μl) was injected to an Ascentis C18 2.7μm 2.1×150mm column. Mobile phases consisted of A (acetonitrile/water(60:40)) and B (isopropanol/acetonitrile (90:10)) in the presence of ammonium formate and formic acid. Lipid species were detected using sMRM in the positive-ion mode reflecting the neutral loss of (RCOO + NH3) for DAG and TAG, the product ion scan of m/z 369 (cholesterol – H_2_O) for CE and FC.

#### Structural elucidation

Structural determination of major PL (PC, PE, PI) chains presented in Table S2 was performed by LC–MS/MS using reversed-phase separation on a Symmetry shield RP8 50 mm × 2.1 mm, 3.5 μm column (Waters Corporation, Milford, MA, USA) as previously described (26) and negative ionization using precursor ion scans of FA chains.

#### Data processing

An in-house developed R script was used to correct for isotopic contribution on MRM signals from HILIC injections. Features with over 80% missing values were discarded, missing values of the remaining features were imputed using the KNN approach on the MetaboAnalyst open source software. Lipid features whose variability exceeded 30% in the quality controls were removed.

Lipid amounts were expressed as mole percent of total lipids, except for the plasma lipid data that were given in nmol/μl of plasma to allow full comparison with previous reports (27, 28). Such relative quantification presents the advantage to overcome any uncertainties relative to the measurement of protein concentrations from different sample type and exclude any bias relative to the impact of adipocyte hypertrophia on protein content.

#### Lipid nomenclature

Lipids are abbreviated as follow: Neutral lipids (NL) – cholesteryl ester (CE), diacylglyceride (DAG), triacylglyceride (TAG); free cholesterol (FC); sphingolipids (SL) – ceramide (Cer), dihydroceramide (DHC), sphingomyelin (SM); Phospholipids (PL) – phosphatitic acid (PA), phosphatidylcholine (PC), phosphatidylethanolamine (PE), phosphatidylglycerol (PG), phosphatidylinositol (PI), phosphatidylserine (PS) and the lyso (L) species (LPC and LPE). Plasmalogen linkages are denoted by p- (ex : PEp).

The side-chain structures are denoted as “carbon chain length:number of double bonds” and are provided for each chain where they could be determined, or as a total number of all carbons and double bonds where individual chains could not be determined.

#### Statistical analysis

Comparison of sample types (plasma, EV subtypes and VAT) and sample groups (lean or obese) were run using paired Wilcoxon, Mann-Whitney-test. Pairing was either sample driven (IEV and sEV from the same mice) or batch driven (lean and obese samples from the same collection date). In order to overcome the batch effects between two studies, we used a Relative difference (RD) calculation strategy, where we calculated the RD between the subject groups within a batch by transforming them into an expression value using MeV software version 4.9 (https://sourceforge.net/projects/mev-tm4/) (25). The outcome of the analysis produces a transformed RD data with log2-fold change values that corresponds to the differences between the given subjects, notified as ‘vs’ (for versus) in the legends. Hierarchical clustering tree (HCL heatmap) were produced from clustered normalized mean-center data using complete linkage over the features using Pearson correlation distance matrix. Features were considered significant when the p-value was below 0.05 after Benjamini-Hochberg correction controlled for false discovery rate (FDR) (29).

## Results

### Lipidomic analysis of *obese* mice plasma reflects common lipid alterations associated with obesity

In order to perform comparative lipidomic analysis between the obese and lean state independent of changes in dietary lipid sources, we investigated the leptin-deficient (*ob*/*ob*) mouse which develops obesity on standard chow diet due to spontaneous hyperphagia, but does not require feeding on a high fat diet. We used a complex lipid profiling method based on LC-ESI^+^-MS/MS analysis that combine HILIC LC separation mode with scheduled multiple reaction monitoring (sMRM) to assess plasma levels of different lipid classes, including phospholipids, sphingolipids and sterols, which have been shown to act as biomarkers or active contributors of obesity-associated metabolic complications (30, 31). Prominent increases in PC, LPC, PI, SM, Cer as well as cholesterol esters and free cholesterol were observed in *ob/ob* mice (Figure 1A-C). These alterations also impacted specifically some other lipid subspecies providing a lipid fingerprint of *ob/ob* mice plasma (Table S3). Our results are in agreement with previous lipidomic analysis performed on *ob/ob* mice plasma that highlighted specific increase in sphingolipids (especially Cer and SM), cholesterol esters and PC subspecies as a hallmark of obesity (27, 28).

**Figure 1 :**
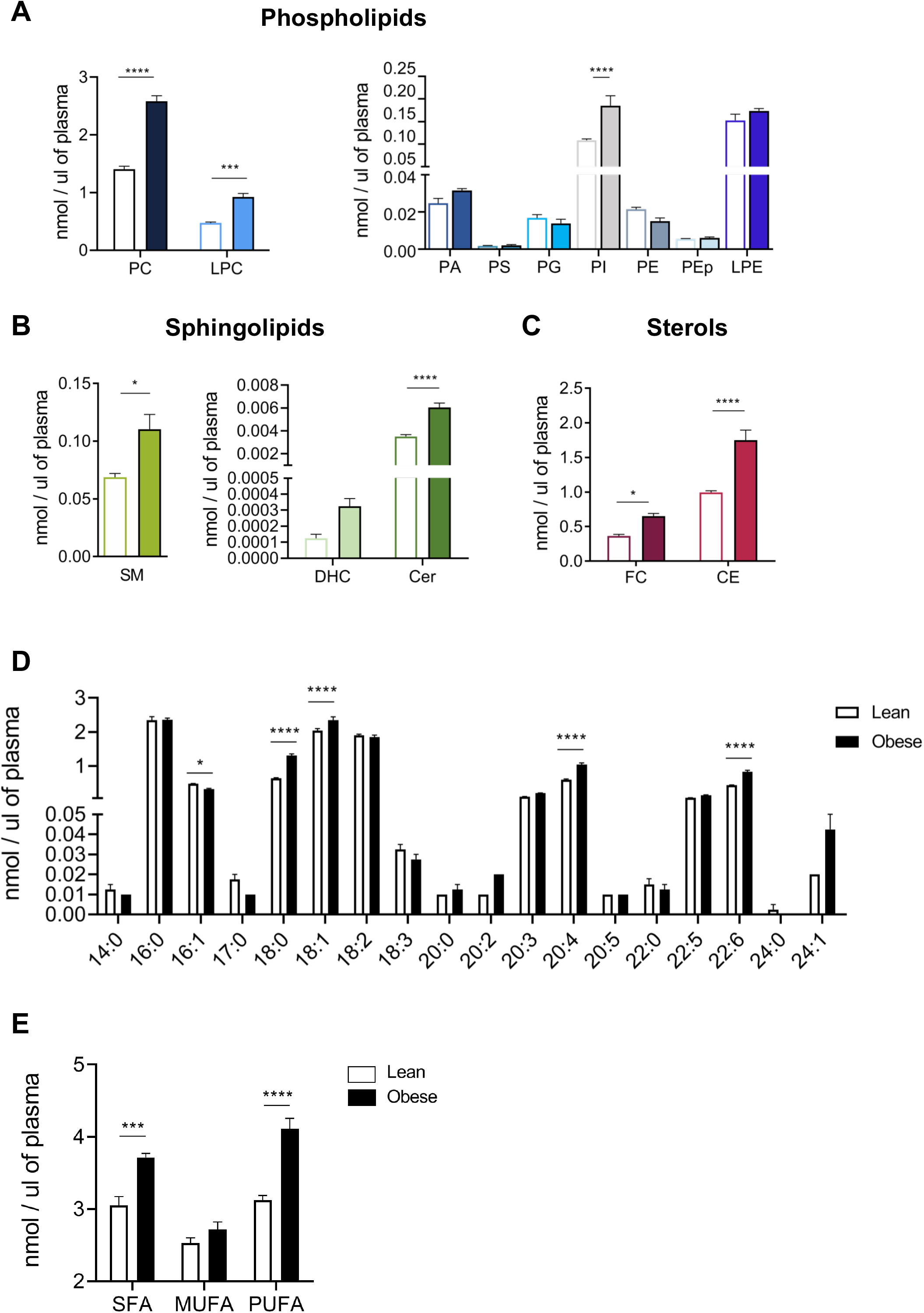
Plasma lipidome in lean and obese mice. **A-C**. Absolute lipid composition of circulating lipid species expressed in nmol/μl plasma. Major lipid families are distinguished : phospholipids (**A**), sphingolipids (**B**) and sterols and glycerolipids (**C**). Minor and major lipids are presented on different graphs for the different families. Void bars are lean conditions, filled bars are obese conditions. **D**. Fatty acid composition of plasma phospholipids based on structural elucidation of phospholipids as presented in Table S2. Lipids were regrouped according to the number of carbon atoms present in their acyl chains. Note that the ‘y axis’ is discontinuous. **E**. Unsaturation profiles of plasma phospholipids. Phospholipids were classified as saturated (SFA), monounsaturated (MUFA) and polyunsaturated (PUFA) lipids based on structural elucidation as presented in Table S2. Lipids were regrouped according to the number of double bonds present in their acyl chains. SFA are fully saturated lipids on both chains, MUFA has at least one or 2 monounsaturated chain and no polyunsaturated chain and PUFA have at least one polyunsaturated chain out of the 2. Results are presented as the mean ± SEM calculated from four independent samples for each metabolic state. Asterisks indicate a significant difference between lean and obese (p-value<0,05*, p<0,01**,p<0,005***, p<0,001****, non-parametric two-way ANOVA test corrected for multiple comparisons by Sidak’s test).

Fatty acid distribution on the most abundant phospholipids classes (PC, PE, PI) was elucidated using negative ion mode LC-MS/MS. Fatty acid distribution of minor phospholipids (PS, PG, PA) was extrapolated from these data and from the literature (32) (see Methods and Table S2). This revealed an increase in total lipids containing 18 carbon atoms including stearic acid (C18:0) and oleic acid (C18:1), at the expense of palmitoleic acid (C16:1) (Figure 1D). Besides, a significant increase in arachidonic acid (20:4), mainly retrieved in plasma under its omega 6 form (32), and in docosahexaenoic acid (22:6), the final product of omega-3 fatty acid elongation and desaturation, was also observed likely reflecting higher uptake of essential fatty acids through diet by hyperphagic *ob/ob* mice (31). By this mean, we noticed that plasma lipids from *ob/ob* mice are overall enriched in saturated fatty acids (SFA) and polyunsaturated fatty acids (PUFA) compared to lean controls (Figure 1E).

Altogether, plasma lipid profiling of *ob/ob* mice confirmed circulating lipid biomarkers of obesity. Significantly altered lipid moieties in obese mice plasma recapitulated global obesity-related changes in human plasma lipidome (30, 33–35), reinforcing the relevance of using the *ob/ob* mice as a preclinical model of obesity.

### Obesity impacts adipose tissue lipidome, paralleling plasma lipid alterations

We next investigated the lipid content of VAT collected from 3-month lean and genetically obese mice, that will be further use to produce VAT-derived EV. We provided a comprehensive profiling of the VAT lipidome including phospholipids, sphingolipids, neutral lipids (TAG, DAG) and cholesterol (FC and CE). A total of 354 lipid species were scanned, among which 265 passed the quality control and were subsequently quantified. Based on calibration with internal standards, data were normalized either to total lipid quantified (including neutral lipids) or to total membranous lipids (SL plus PL) and expressed as mol% to investigate the relative changes of AT lipid composition between lean and obese mice.

As expected, the vast majority of lipids in VAT samples were TAG – the lipid form of AT energy stores - reaching 90% of all identified lipids regardless of metabolic condition (Figure 2A). DAG, which is associated with TAG turnover, represented only minor neutral lipid stores (<8%), whereas only traces of CE were detected (0.01%). Besides neutral lipids, other lipids accounted for only 3% of total lipids and mainly included phospholipids and free cholesterol (FC), as well as sphingolipids in smaller proportions. Among phospholipids, PC and PE were the most abundant classes thereby providing the majority of membrane lipids within adipose cell. To explore how obesity context affected the sphingophospholipidome of VAT, we assessed the relative differences of lipid species between lean and obese VAT (Figure 2B). By this mean, we observed significant enrichment of obese VAT in total PG and PI, reflecting respectively higher content of the predominant PG(34:1) and PI(38:4) lipid species in VAT (Figure S1). Conversely, total LPE and PE plasmalogens (PEp) were significantly decreased in *ob/ob* VAT (Figure 2B).

**Figure 2 :**
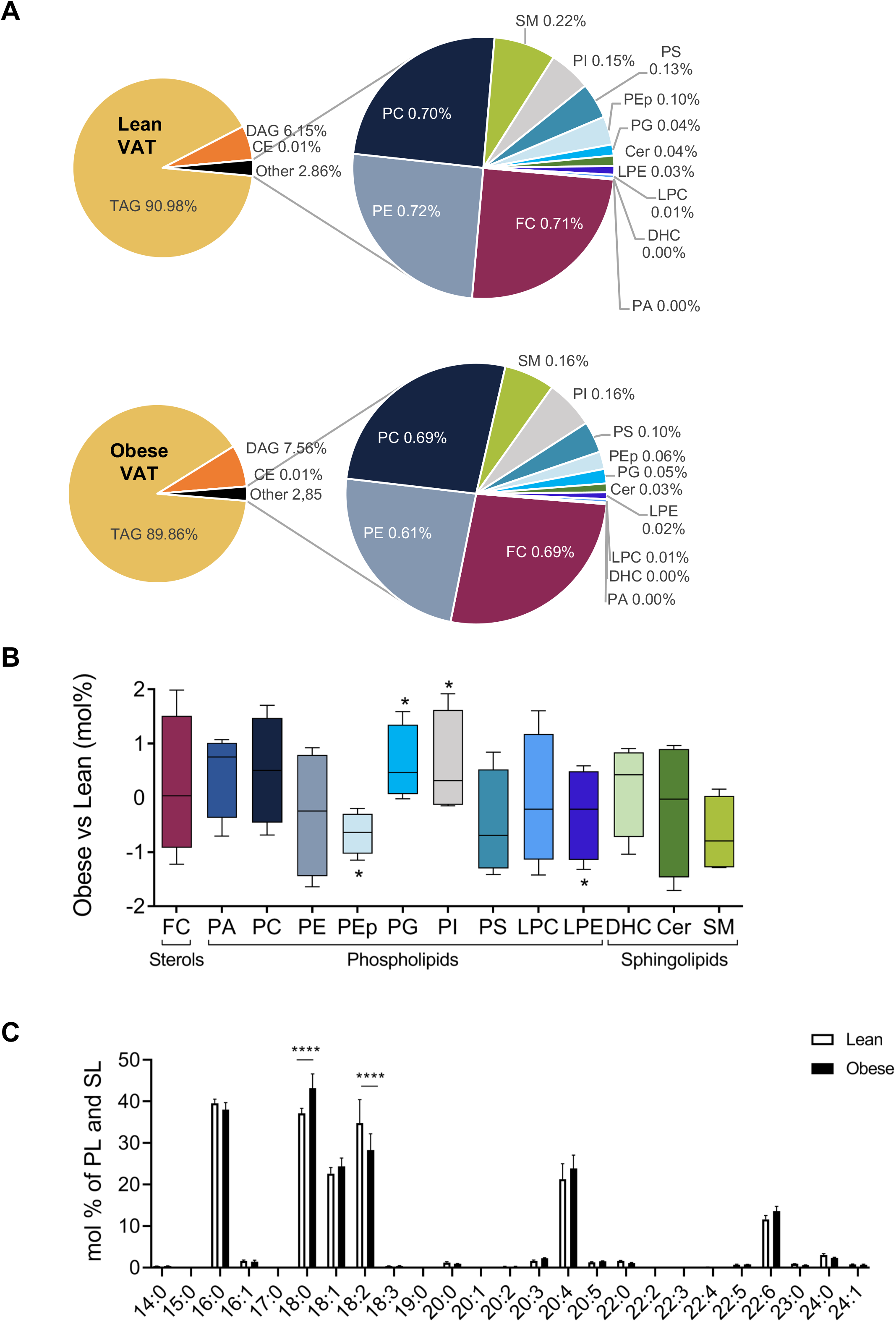
VAT lipidome in lean and obese mice. **A.** Pie diagram showing the relative lipid classes composition of lean and obese VAT. Besides neutral lipids, other lipids correspond to sphingolipids and phospholipids which are detailed in a sub-pie diagram. Data presented correspond to the mean amount (in mol %) respectively in lean VAT (left pie charts) and obese VAT (right pie charts) from five independent samples for each metabolic state. **B.** Relative membrane lipid classes composition of VAT membrane lipids between the obese and lean state. The panel bar plots represent the difference between the mean of obese and lean in the VAT calculated from four independent samples for each metabolic state. Asterisks indicate the lipid species that differ significantly between the two metabolic states (see methods for statistical analysis, p-value<0,05). **C-D**. Total acyl chain length **(C)** and unsaturation profile **(D)** elucidated for sphingolipids and phospholipids of VAT from lean and obese mice. Lipids were grouped according to the number of carbon atoms (**C**) and the number of double bonds (**D**) present in their acyl chains. Complete acid chain length profiles are presented in Table S3 based on structural elucidation performed according to Table S2. Results are presented as the mean ± SEM calculated from four independent samples for each metabolic state. Asterisks indicate a significant difference between lean and obese (p-value<0,05*, p<0,01**,p<0,005***, p<0,001****, non-parametric two-way ANOVA test corrected for multiple comparisons by Sidak’s test).

Structural elucidation of membranous lipids on VAT samples revealed a significant increase in C18:0 acyl chains containing lipids, at the expense of C18:2 containing lipids, and a trend to an elevation of polyunsaturated lipids (number of total double bounds>4) (Figure 2C). These lipid enrichments parallel lipid disturbances observed in *ob/ob* mice plasma (see Figure 1D-E), in agreement with the primary lipid storage function of VAT.

Overall, we demonstrated distinct lipidomic profile between lean and obese VAT mirroring the changes in plasma lipidome.

### AdEV subtypes secretion is enhanced with obesity

In order to isolate AdEV subtypes from secreted VAT explants collected from lean and obese mice, we implemented a culture system using Clinicell^®^ cassettes, allowing optimal gaz/air exchanges ensuring the full viability of VAT explants (23) (Figure 3A). EV isolation was done from 48h-serum free VAT explant conditioned media using differential (ultra)centrifugation (respectively 13 000 x*g* for IEV and 100 000 x*g* for sEV). TEM imaging on VAT-derived AdEV confirmed the successful isolation of large vesicles surrounded by a kind of matrix layer, by comparison to smaller electron-dense vesicles homogenous in size (Figure 3B). Larger size for lean IEV compared to lean sEV was quantified by NTA measurements, whereas size of AdEV subtypes isolated from obese VAT are very dispersed rendering size differences between obese sEV and obese IEV unsignificant (Figure 3C). Obese IEV concentrations were moreover significantly higher than lean IEV (Figure 3D). A trend towards higher concentrations of obese sEV by comparison to lean sEV was also observed, in agreement with the highly concentrated AdEV productions previously described from high-fat diet mice (5, 14). Biochemical analysis of AdEV subpopulations demonstrated specific enrichment of tetraspanins (CD9, CD63) in sEV by comparison to IEV or secreting VAT (Figure 3E). Conversely, flottilin-2 or Grp94, that we previously identified as specific markers of 3T3-L1 adipocyte-derived IEV (19), were specifically enriched VAT-derived IEV (Figure 3E). No perilipin-1 signal was retrieved in any AdEV preparations suggesting absence of lipid droplet containing vesicles (Figure 3E).

**Figure 3 :**
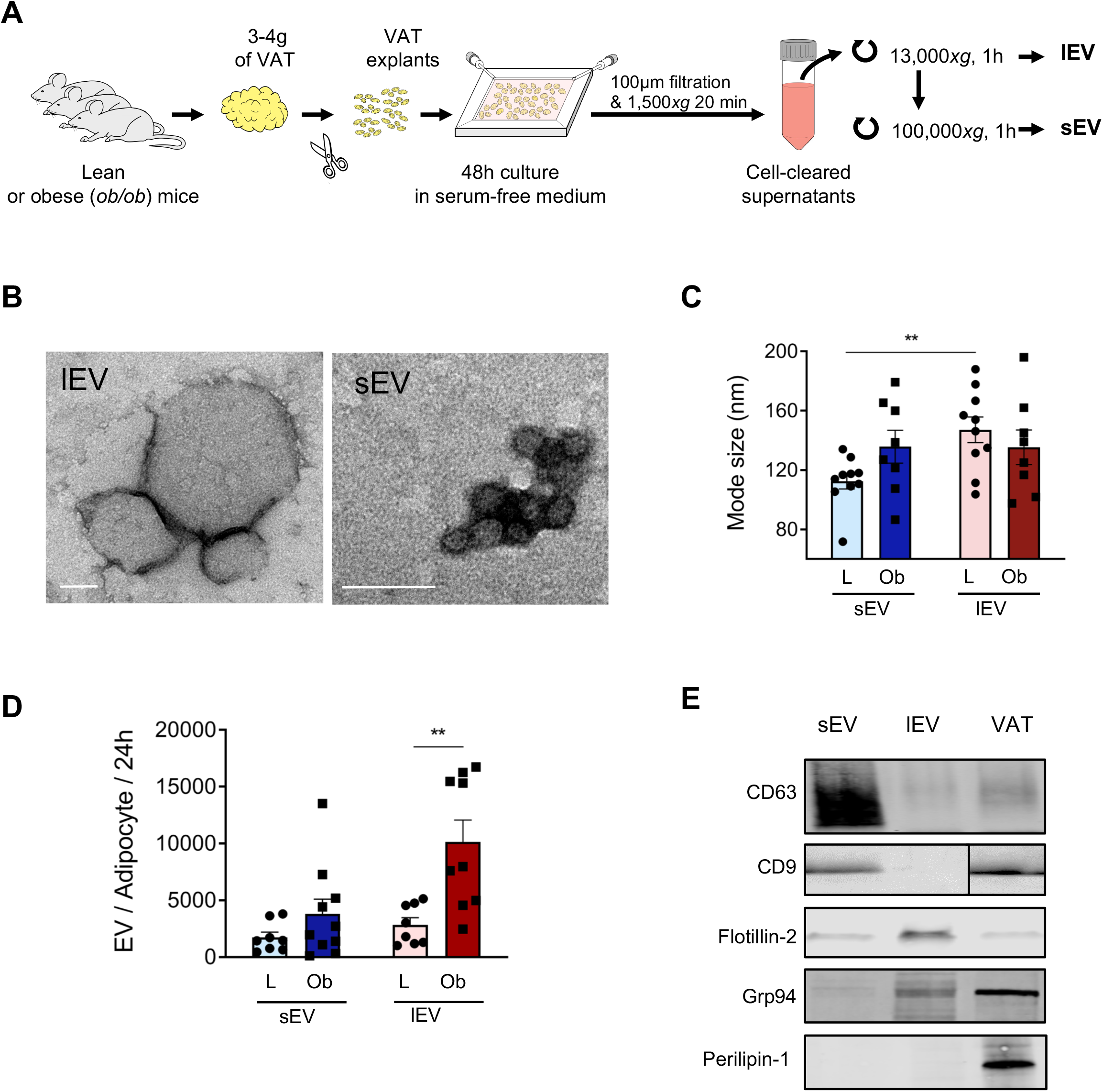
AdEV isolation and characterization. **A.** Schematic representation of VAT-derived EV isolation leading to AdEV subtype isolation, namely IEV and sEV **B.** Transmission electron microscopy images of IEV (upper panel) and sEV (lower panel) pellets. Scale bar : 100nm. **C-D**. Nanoparticle tracking analysis (NTA) of IEV and sEV populations isolated from VAT-derived conditioned media. IEV (13,000 × g pellet) and sEV (100,000 × g pellet) were collected from VAT conditioned media and resuspended in NaCl. EV mode size **(C)** and EV concentrations (expressed as EV numbers secreted by adipocyte cell) **(D)** were analyzed. No impact of obesity on EV size was observed **(C)**. Abundant EV secretion per adipocyte was observed for IEV and sEV, which is moreover significantly increased by obesity **(D)**. In total, 8 to 10 independent IEV and sEV preparations per metabolic state were analyzed by NTA. Results are presented as mean ± SEM (p-value<0,05*, p<0,01**, Wilcoxon matched pairs rank test for **C**, non-parametric two-way ANOVA test corrected for multiple comparisons by Sidak’s test for **D**). **E**. Western blot analysis of different EV markers in sEV and IEV preparations. Eight micrograms of VAT explant, IEV or sEV samples were migrated on a SDS-PAGE gel and analyzed by immunoblotting for the tetraspanins CD63, CD9, flotillin-2, Grp94 and perilipin-1. One representative blot for each protein is presented.

### Enrichment of various lipid species in AdEV compared to source VAT illustrates specific EV lipid sorting

Qualitative comparisons between AdEV and VAT lipidomic datasets by principal component analysis (PCA) showed that AdEV segregated from source VAT along the first principal component (PC1) which explained nearly 70% of the variance (Figure 4A). AdEV or VAT separated along PC2 according to the metabolic context (Figure 4A). This highlighted active specific sorting of lipids by VAT-derived EV secretion. Of note, sEV and IEV overlapped in the lean or in the obese context illustrating that AdEV lipid composition was mainly driven by the metabolic context rather than by AdEV subcellular origin.

**Figure 4 :**
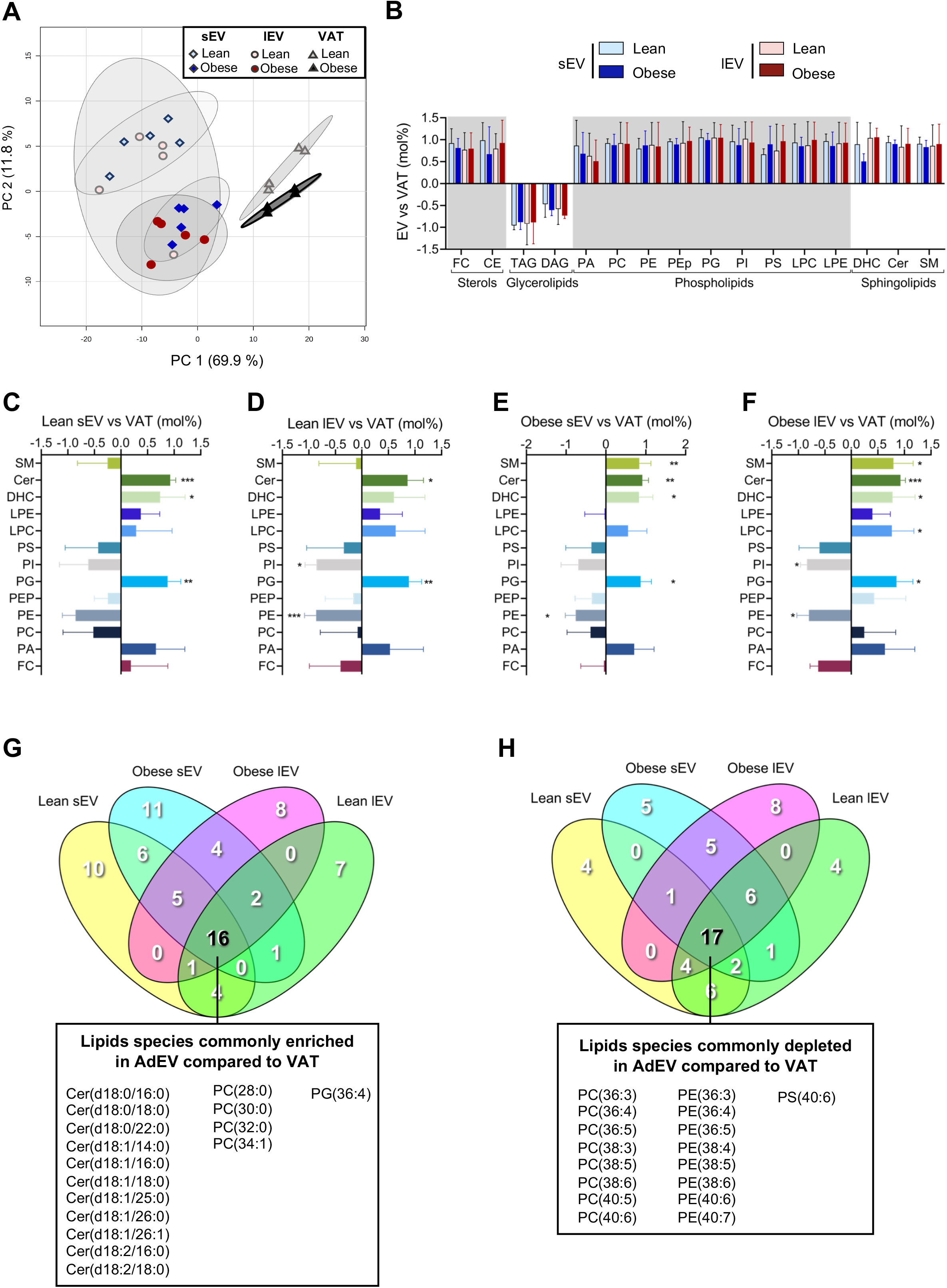
AdEV relative lipid composition compared to VAT. **A.** Principal Component Analysis (PCA) of VAT and EV lipidomes in lean and obese metabolic state. PC1 segregated EV from VAT whereas PC2 segregated samples according to lean or obese metabolic state. sEV and IEV co-segregated in the lean or obese state highlighting the metabolic state as an important driver of EV lipid content. **B.** Relative lipid class composition of VAT and VAT-derived EV in the lean and obesity context. Bar plots represent the relative differences (in mol%) between AdEV versus VAT for the respective lipid class analyzed. Significant enrichment of AdEV was observed for sterols, phospholipids and sphingolipids whereas AdEV were significantly depleted in glycerolipids (see methods for statistical analysis, p-value<0,05 for all lipid class analyzed, no asterisks indicated in the panel for better lisibility). **C-F**. Relative membranous lipid class composition of sEV and IEV by comparison to source VAT. Relative differences (in mol%) between AdEV and VAT for the respective lipid class analyzed were calculated in the lean (**C-D**) and obesity context (**E-F**), by distinguishing sEV (**C ; E**) from IEV (**D ; F**). All results in **(B-F)** are presented as the mean ± SEM, calculated from four independent samples for each EV subtype and related source VAT for each metabolic state (see methods for statistical analysis, p-value<0,05*, p<0,01**,p<0,005***). **G-H**. Venn diagrams depicting the lipid features that share commonness and uniqueness in their expression in lean sEV, lean lEV, obese sEV and obese IEV against the VAT. Lipid features that are enriched (**G**) or depleted (**H**) are presented. Detailed list of lipid species enriched or depleted in the different subsets are provided in Table S4 and Table S5, respectively.

Global lipid composition changes between AdEV and VAT mainly related to VAT enrichment in glycerolipids (TAG and DAG) whereas phospholipids and cholesterol were abundantly retrieved in AdEV subtypes, in agreement with their membranous origin (Figure 4B). Thus, data were subsequently expressed as mole % of total PL and SL to focus on membrane lipids. This revealed a significant enrichment in Cer, DHC and PG lipid classes for both sEV and IEV versus lean VAT (Figure 4C-F). Conversely, PI and PE were significantly depleted from lean lEV, with a similar trend observed for lean sEV, by comparison to lean VAT phospholipidome (Figure 4C-D). Similar depletions were observed for obese sEV and IEV compared to source obese VAT (Figure 4E-F). Noticeably, AdEV enrichment in SM by comparison to source VAT was observed in obese samples (Figure 4E-F).

We next investigated specific lipid species either significantly enriched (Figure 4G and Table S4) or depleted (Figure 4F and Table S5) in AdEV subtypes by comparison to source VAT. Sixteen lipid species were found commonly enriched in AdEV, independently of AdEV subtype or metabolic context, which included 11 Cer and 4 PC subspecies as well as PG(36:4) (Figure 4G). By contrast, 17 lipid species commonly depleted in AdEV compared to VAT were identified that included 8 PE subspecies, 8 PC subspecies and PS(40:6) (Figure 4H).

Altogether, compared to source VAT, overall AdEV lipid composition pointed to Cer, DHC and PG enrichment and selective SM accumulation in obesity. Lipid species commonly enriched in AdEV, independently of the pathophysiological state, moreover revealed the selective lipid sorting of Cer subspecies, including DHC, and PG(36:4) as well as some specific PC lipid subspecies. Overall, these lipids represent tracers of AdEV lipid transport and relevant candidates for lipid-associated AdEV mediated signaling in target cells.

### Lipid fingerprints differentiate VAT-derived EV subtypes

We next studied how AdEV lipidome is influenced by EV subtype. Comparison of the relative distribution of all lipid classes screened between IEV and sEV highlighted a striking enrichment of TAG in large vesicles by comparison to smaller ones (Figure 5A-B), but no significant changes were observed in CE or DAG content of IEV and sEV (Figure 5B). Alternatively, sEV displayed a specific FC enrichment compared to lEV, whatever the metabolic status considered (Figure 5C). Among phospholipids, we demonstrated enrichment of IEV in total PC and total LPC, two major structural membrane lipids, as well as in total DHC (Figure 5D-E). At individual species resolution, lean IEV were specifically enriched in short LPC and 34:1 lipid subspecies (PC, PE, PG and SM) and depleted in long polyunsaturated phospholipids over almost all classes (Figure S2).

**Figure 5 :**
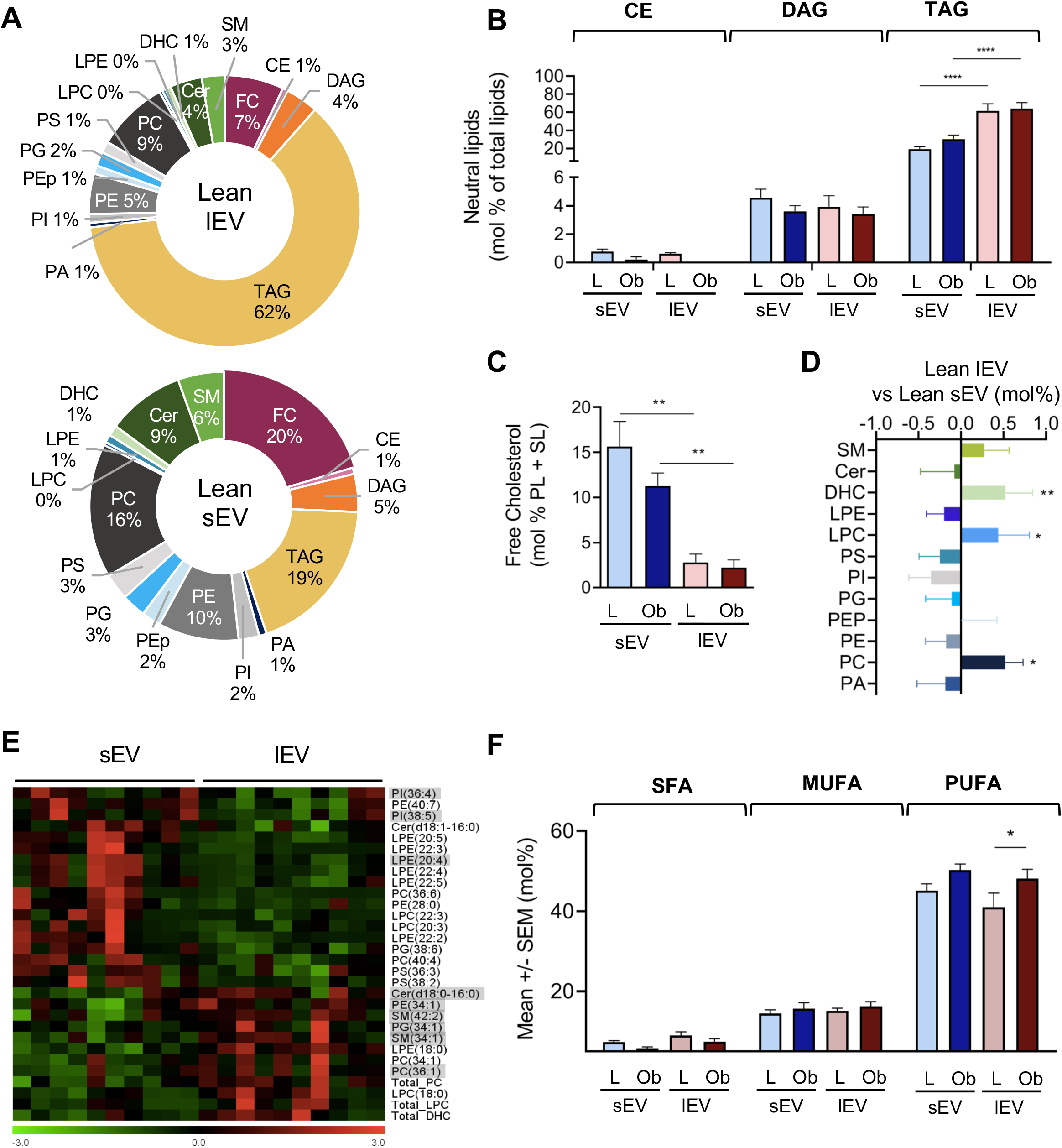
AdEV subtypes relative lipid composition in the lean and obese context. **A.** Pie diagram showing the relative lipid classes composition of AdEV subtypes isolated from lean VAT. A striking TAG enrichment is observed for lean lEV, at the expense of free cholesterol (FC), phosphatidylcholine (PC) and ceramides (Cer) by contrast enriched in lean sEV. Data presented correspond to the mean amount (in mol %) respectively in lean IEV (upper pie chart) and lean sEV (lower pie chart) from five independent samples for each AdEV subtypes. **B.** Relative neutral lipid composition of AdEV subtypes isolated from lean and obese VAT conditioned media. Specific enrichment of total TAG (expressed in mol% of total lipids quantified) is observed in IEV compared to sEV, whatever the metabolic state considered. Data are presented as mean ± SEM, ****p < 0.001 (non-parametric two-way ANOVA test corrected for multiple comparisons by Tukey’s test). **C.** Relative free cholesterol (FC) composition of AdEV subtypes isolated from lean and obese VAT conditioned media. Small EV displayed specific FC enrichment (expressed as mol% of membranous lipids) in the lean and obese states compared to lEV. Data are presented as mean ± SEM, **p < 0.01 (non-parametric two-way ANOVA test corrected for multiple comparisons by Tukey’s test). **D.** Relative membranous lipid class composition of AdEV subtypes. Relative differences (in mol%) between lean IEV versus lean sEV for the respective lipid class analyzed were calculated. Asterisks indicate the species that differ significantly between IEV and sEV (see methods for statistical analysis, p-value<0,05*, p<0,01**). **E.** Heatmaps of AdEV subtypes lipid fingerprints in the lean context. Lipid subspecies that significantly (p <0.05) differentiated the lean sEV and the lean IEV are plotted in a hierarchical clustering tree (HCL). In red are lipid species that are enriched, and in green are those that are depleted. Major lipid species for each lipid class are underlined in grey. **F.** Unsaturation profiles of AdEV phospholipids. Phospholipids were classified as saturated (SFA), monounsaturated (MUFA) and polyunsaturated (PUFA) lipids based on structural elucidation as presented in Table S2. Data are presented as mean ± SEM from 10 lean or 8 obese independent samples, *p < 0.05 (non-parametric two-way ANOVA test corrected for multiple comparisons by Sidak’s test).

Obesity impacted similarly IEV and sEV by favoring total PI and PC and by increasing the amount of AdEV-associated major PG(34:1) (Figure S3-S4), partly mirroring obesity-associated lipid changes previously observed in VAT (Figure 2B). Significant total SM and/or SM subspecies enrichment was moreover observed for all AdEV subtypes isolated from obese VAT by comparison to AdEV derived from lean animals, whereas Cer-associated AdEV were by contrast decreased in obesity context (Figure S3 and S4C-D). These similar lipid profiles changes in AdEV subtypes are in agreement with the lack of size differences observed between IEV and sEV when isolated from obese VAT (Figure 3C). These data therefore illustrated that AdEV lipidome is strongly influenced by source VAT, and that isolation of EV subtypes based on vesicle size, for lipidomic studies, is irrelevant in the context of obesity.

Finally, lipid structural elucidation allowed us to pinpoint common altered AdEV lipid species alterations with obesity. We therefore highlighted enrichment of AdEV subtypes isolated from obese VAT in arachidonic acid (20:4)-containing species including 36:4 (PI and PE), 38:4 (PI, PC, PE, PEp) and 38:5 (PI and PE) species (Figure S3 and S4E-F). We moreover demonstrated significant enrichment of 18:1-containing species as for LPC(18:1), PG(34:1), PC(34:1) and PC(36:1) in AdEV subtypes derived from obese VAT, at the expense of short and saturated palmitic-containing species as for 32:0 and 34:0 phospholipids (Figures S3 and S4). This translated into a trend for decreased total saturated fatty acid (SFA) in obese IEV and sEV, favoring obese VAT-derived AdEV PUFA enrichment (Figure 5F). Finally, among PE plasmalogens, that are known to display antioxidant properties, we found five species commonly depleted in obese IEV and sEV (PE(16:0p/18:2) PE(18:0p/18:1) PE(18:0p/18:2) PE(18:0p/20:4) PE(18:0p/20:5)) (Figure S3), reflecting significant PEp decrease in source obese VAT (Figure 2B) which may favor adipose oxidative stress.

## Discussion

This study aimed to define lipid fingerprints of AdEV subtypes isolated from lean and genetically obese (*ob/ob*) mice in order to determine the signaling capacity of AdEV-transported lipid species in obesity-related complications. We provide here comprehensive lipid maps revealing specific adipose EV lipid sorting when compared to secreting VAT. We demonstrated that the EV lipidome is highly influenced by the pathophysiological state of VAT and identified some specific AdEV lipid classes and species closely related to obesity status. Notably, we suggest that AdEV lipid subspecies enrichment may contribute to the development of metabolic dysfunctions associated with obesity.

Our comparative analysis between source VAT and secreted AdEV revealed distinct lipidomic profiles between both sample types. As expected, AdEV are preferentially composed of membranous lipids in line with their biogenesis. According to PCA, AdEV lipid composition appeared to be much more influenced by the metabolic status (lean or obese) rather than the EV subtype (IEV or sEV). Thus, although major differences in protein contents were reported among EV subtypes, we highlight here an overall stability of the lipid composition of large versus small EV. Moreover, EV subtype distinction based on EV size (large or small) appeared of minor relevance in the obese context. Nonetheless, separation of IEV from sEV from lean VAT allowed us to pinpoint specific lipid features of each EV subtype that might be of particular relevance, especially cholesterol enrichment in sEV, a feature that we already highlighted in sEV isolated from 3T3-L1 adipocytes (19). TAG were found enriched in large EV preparations. These apolar lipids are likely to be localized within the core of the vesicles as reported in a recent study identifying adipocyte-derived EV as neutral lipid-filled vesicles (13). However, it is noteworthy that TAG-enrichment of large vesicles is limited enough to allow particle sedimentation by centrifugation. The absence of perilipin-1 (a specific marker of adipose lipid droplets) argues against EV-based extrusion of lipid droplet organelles, and would rather suggest a TAG-IEV loading which may be interconnected to lysosome-mediated lipid droplet degradation.

Besides these specific EV subtype lipid traits, some EV-lipid enrichments are common to both EV subtypes. We therefore revealed specific sphingolipid (SM, Cer, DHC), LPC and PG AdEV enrichment. Increased relative proportion of cholesterol, Cer and SM have already been measured in different types of EV, and would be related to the involvement of these lipids in EV biogenesis (10, 36, 37). This type of lipid enrichment may contribute to increase the rigidity of sEV (38), provide higher sEV membrane order degree (39) and increase sEV resistance to non-ionic detergents (40) as it is the case for lipid raft membrane microdomains. Such lipid composition certainly confers an advantage for the stability of these EV in biological fluids and/or their binding or uptake by recipient cells (41).

Conversely, we observed relative decrease of PE and PI in AdEV compared to source VAT, particularly in obese lEV. Such depletion has been previously observed in EV isolated from different cell sources, and was partially compensated by PS sEV enrichment (10). However, it remains unclear how this can impact EV structure since PE and PI are in minority compared to PC EV proportions, whose EV content is moreover enhanced with obesity.

Obesity-associated circulating lipid alterations translated at the level of organs, particularly AT or liver. AT lipidome of diet-induced obese mice is featured by a significant increase in longer and more unsaturated TAG and PL species and characterized by accumulation of lipids made of acyl chains containing 18 carbons (42). Similarly, lipidomic analysis performed on AT from twin pairs discordant for obesity showed that membrane lipids containing longer and more unsaturated fatty acids were more abundant in the obese by comparison to the lean individuals (34). We indeed observed elevated PUFA and C:18 fatty acid increase in plasma, VAT and AdEV suggesting overall lipid equilibrium between plasma, VAT and VAT-derived AdEV. Besides these overall lipid alterations, numerous studies have pinpointed sphingomyelin metabolites as signaling molecules of pathological biological events related to metabolic dysfunction and identified significant correlations between circulating Cer and DHC as biomarkers of the development of T2D in obese patients (43, 44). In accordance with this hypothesis, we also found elevated plasma Cer levels in obese animals. However, a surprising finding from our study was the relative decrease in ceramides in AdEV derived from obese mice, despite AdEV Cer enrichment with regard to source VAT. Nonetheless, our data are in line with the previously reported increased ceramidases activities and the associated decreased ceramide levels in VAT from *ob/ob* mice (27). Whereas adipose ceramides have been shown to be critical for driving AT remodeling and controlling whole-body energy expenditure and nutrient metabolism (45), our results suggest that the source of elevated circulating ceramides in obesity is likely to be of hepatic origin. Further studies are warranted to establish whether the defect of sphingolipid metabolism in AT observed in *ob/ob* mice is also present in humans.

The study identified AdEV lipid mediators as potential signaling molecules to explain the pathophysiological responses mediated by AdEV isolated from obese VAT (5, 6). As previously demonstrated for proteins and genetic material, AdEV displayed specific lipid fingerprints indicative of oriented EV lipid sorting. We highlighted some specific lipid enrichment of particular relevance in obese VAT-derived AdEV, which can relay metabolic alterations. Particularly, we demonstrated enrichment of AdEV subtypes isolated from obese VAT in arachidonic acid (20:4)-containing species, which are known to mediate inflammatory signaling notably through the production of prostaglandins, thromboxanes, leukotrienes and lipoxins. Their production is mainly mediated through the action of PLA2, which allows the release of different PUFA from membrane PL.

Therefore, these signaling lipids may be either carried from the parental cells or directly generated within EV, since exosomal activable PLA2 have been detected in exosomes from the mast cell line RBL-2H3 (46). PLA2 from the extracellular milieu may also act on phospholipid EV, as previously described for secreted PLA2-IIA present in inflammatory fluids which acted in concert with EV-associated platelet-type 12-lipoxygenase to generate autonomously 12(S)-hydroxyeicosatetranoic acid within EV (47). EV-associated PLA2 may also contribute to raise EV-LPC content which may serve as a substrate for autotaxin-bound EV therefore contributing to raise the bioactive lipid lysophosphatidic acid (LPA) (48). Knowing the lipogenic, anti-lipolytic and inflammatory role of PLA2-downstream mediators in obesity (49), our data confirm the potential interest of considering EV eicosanoid content to be informative of the metabolic status (i.e. healthy vs obese) (12).

PG subspecies were also found enriched in AdEV from obese mice, reflecting PG lipid class relative accumulation in obese VAT. PG are specific mitochondrial phospholipids, precursors of cardiolipins. Previous studies revealed that PG serum concentration in obese patients was the lipid class (among other blood phospholipids) that was predominantly and positively associated with body mass index and with AT inflammation (50). Importantly, serum PG levels sharply declined after metabolic improvement following weight loss either induced by nutritional intervention or bariatric surgery in patients with obesity (51). PG are poorly investigated in most of lipidomics studies, but cardiolipins have been shown to be markedly enriched in sEV from different cell sources (10). Whereas PG can act as lipid mediators notably favoring adipose lipid storage (50), PG enrichment might also be related to the presence of mitochondria within EV, recently defined as mitovesicles (52). A recent study evidenced intercellular mitochondria transfer between adipocytes and macrophages in VAT as a mechanism of immunometabolic crosstalk that regulates metabolic homeostasis and which is impaired in obesity (53). Whether PG AdEV content reflects this mitochondrial extrusion will be need further investigations.

Finally, we found a significant depletion of PE plasmalogens (PEp) in AdEV isolated from *ob/ob* animals as well as in obese VAT. Although these vinyl ether-bound lipids are widespread in all tissues and can represent up to 18% of the total phospholipid mass in humans, their physiological function remains poorly understood (54). PEp have been previously found enriched in EV from platelets (55) and constitute more than half of nematode EV lipid content increasing EV membrane rigidity (56). Interestingly, external addition of an ether lipid precursor to human prostate cancer PC-3 cells to increase cellular ether lipids was found to be associated with changes in the release and composition of exosomes (57). Plasmalogens have been shown to be involved in membrane trafficking and cell signaling, and display some cellular antioxidants properties (54). Atomistic molecular dynamics simulations demonstrated that PEp contribute to form more compressed, thicker and rigid lipid bilayers (58). Future studies are needed to investigate how EV-associated PEp influence their biophysical and biological properties.

To summarize, we identified specific lipid fingerprints for plasma, VAT and AdEV that are informative of the metabolic status and revealed the potential signaling capacity of lipid species transported by AdEV in obesity-related metabolic complications. These findings open some interesting clinical perspective to develop new biomarkers and or drug targets in the obesity context.

## Abbreviation

AdEV: adipose extracellular vesicles
AT: adipose tissue
EV: extracellular vesicles
HOMA-IR: homeostasic model assessment of insulin-resistance
IR: insulin-resistance
lEV: large extracellular vesicles
sEV: small extracellular vesicles
T2D: type 2 diabetes
VAT: visceral adipose tissue

## Data availability statement

Data are available on request.

## Acknowledgments

SLL’ financial supports are Société Francophone du Diabète, la fondation d’entreprise Genavie, INSERM. AB is granted by a PhD allocation from INSERM/Région Pays de la Loire. We thank the SCIAM facility, especially F.Manero, for technical assistance for electron microscopy imaging.

## Author contributions

Methodology, AB, GH, MP, ID, ML, SLL ; Experimentation and analysis, AB, GH, MP, SD, ML, SLL ; writing-original draft preparation, AB, MP, ML, SLL ; review and edition of the manuscript, AB, GH, MP, SD, ID, BC, ML and SLL; conceptualization, AB, SLL; supervision, project administration and funding acquisition, SLL.

All authors have read and agreed to the published version of the manuscript.

## Supplemental Figure legends

**Figure S1 : Detailed phospholipid subpecies in VAT significantly impacted by obesity**

Total LPE, PI, PG and PEp are the phospholipid classes significantly impacted by the obese metabolic state in VAT (see Figure 2B). For each of these phospholipid class, significant changes in lipid subspecies are presented as the relative difference between obese and lean VAT (in mol%) (left panel). Date are presented as the mean ± SEM calculated from four independent samples for each metabolic state. Asterisks indicate a significant difference between lean and obese (p-value<0,05*, p<0,01**,p<0,005***, p<0,001****, see methods for statistical analysis).

Relative abundance of each lipid subspecies detected within each phospholipid class is presented as sidebar color-coded scale (LPE, purple ; PI, black ; blue, PG ; Pep, grey). Relative proportions of each phospholipid subspecies for a given phospholipid class is visually represented by color-gradient, the darkest represent the most abundant subspecies quantified whereas the clearest correspond the less represented ones. Phospholipid subspecies distribution is presented in absolute lipid quantified relative to VAT protein (pmole lipid/mg protein).

**Figure S2: Differential expression of lipids between AdEV subtypes isolated from lean VAT**

Fold change plots of individual lipids in the order of growing chain length in each lipid subclass differentiating lean IEV and lean sEV. Significant lipid species are colored, while non-significant are in grey. Significant threshold: p <0.05.

**Figure S3 : Heatmaps of AdEV subtypes lipid fingerprints in the lean and obese context**

Lipid subspecies that significantly (p <0.05) differentiated the lean and obese metabolic state are plotted in a hierarchical clustering tree (HCL). In red are lipid species that are enriched, and in green are those that are depleted. HCL tree of IEV and sEV are shown in Fig S3A and S3B respectively.

**Figure S4 : Differential expression of lipids retrieved in AdEV subtypes in the lean and obese context**

Fold change plots of individual lipids retrieved in AdEV subtypes, in the order of growing chain length in each lipid subclass, differentiating lean and obese metabolic state. Significant lipid species are colored, while non-significant are in grey. Significant threshold: p <0.05.

## Supplemental Table legends

**Table 1 : Phenotypic characteristics of obese mice and lean control mice**

3 month-old male mice (n= 5-10 for each genotype) were weighed as well as respective organ weight at sacrifice (VAT, SAT, Liver and quadriceps muscles). Blood glucose concentrations and plasma insulin concentrations were measured on the same animals after a 6h fast. Data are presented as mean ± sem, \**p*<0.05, \*\*\**p*<0.001, \*\*\*\**p*<0.001 following student.t tests.

VAT, Visceral Adipose Tissue ; SAT, Subcutaneous Adipose Tissue

**Table S2 : Structural elucidation**

**Table S3 : Plasma lipid fingerprint of lean and obese mice**

Significant lipid changes are expressed as log2 fold change of lipid concentrations in obese mice compared with the respective control mice.

**Table S4 : Detailed lipid species significantly enriched in AdEV compared to source VAT corresponding to Venn Diagram presented in Figure 4G**

**Table S5 : Detailed lipid species significantly depleted in AdEV compared to source VAT corresponding to Venn Diagram presented in Figure 4H**

