## Supplemental files for "Lipidomic analysis of adipose-derived extracellular vesicles reveals their potential as lipid mediators of obesity-associated metabolic complications"

### Figure S1

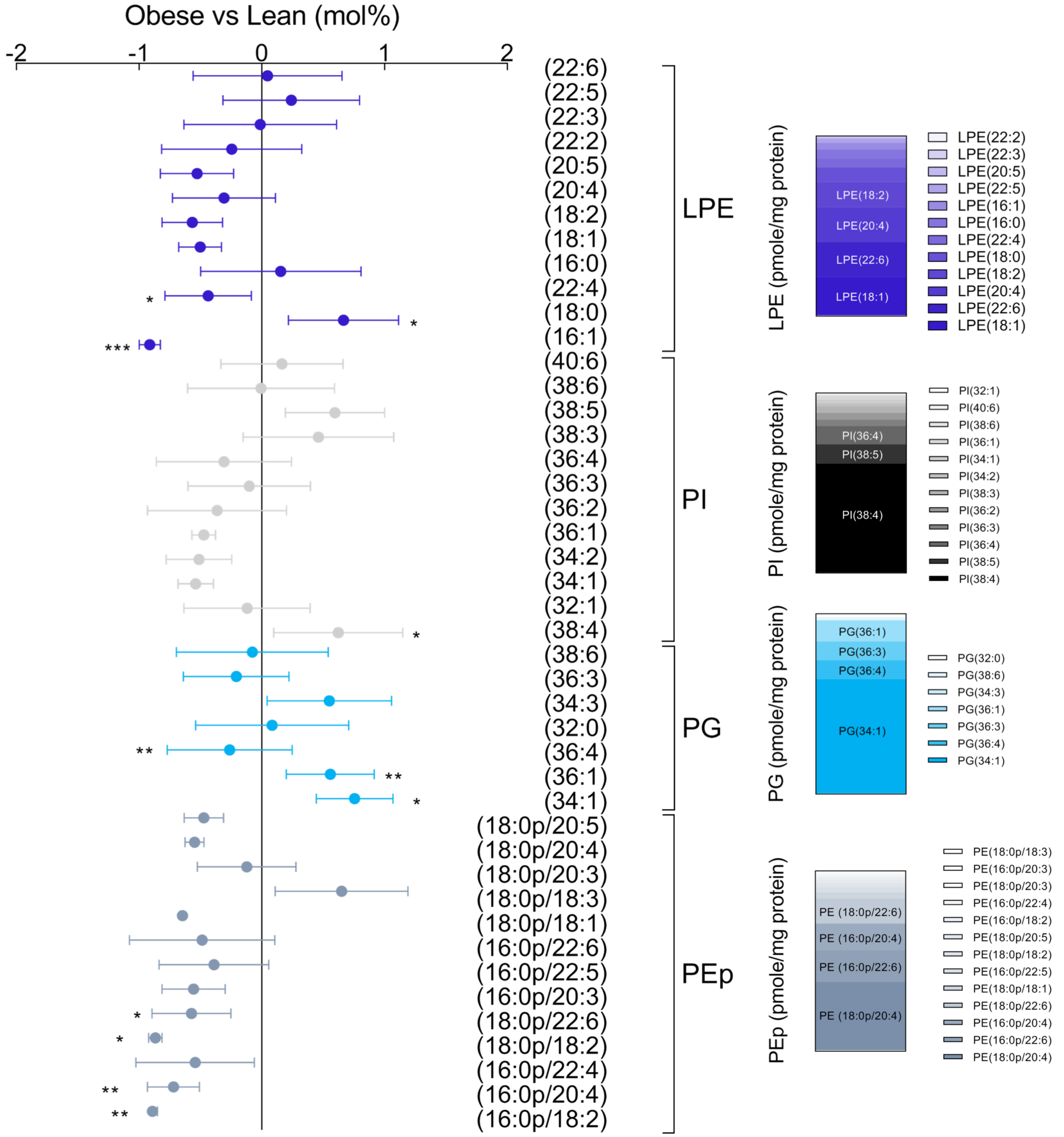
Obese vs Lean (mol%)
-2
-1
0
1
2
(22:6)
(22:5)
(22:3)
(22:2)
(20:5)
(20:4)
(18:2)
(18:1)
(16:0)
(22:4)
(18:0)
(16:1)
LPE
(40:6)
(38:6)
(38:5)
(38:3)
(36:4)
(36:3)
(36:2)
(36:1)
(34:2)
(34:1)
(32:1)
(38:4)
PI
(38:6)
(36:3)
(34:3)
(32:0)
(36:4)
(36:1)
(34:1)
PG
(18:0p/20:5)
(18:0p/20:4)
(18:0p/20:3)
(18:0p/18:3)
(18:0p/18:1)
(16:0p/22:6)
(16:0p/22:5)
(16:0p/20:3)
(18:0p/22:6)
(18:0p/18:2)
(16:0p/22:4)
(16:0p/20:4)
(16:0p/18:2)
PEp
*
*
***
*
**
**
*
*
*
**
**
LPE (pmole/mg protein)
LPE(18:2)
LPE(20:4)
LPE(22:6)
LPE(18:1)
LPE(22:2)
LPE(22:3)
LPE(20:5)
LPE(22:5)
LPE(16:1)
LPE(16:0)
LPE(22:4)
LPE(18:0)
LPE(18:2)
LPE(20:4)
LPE(22:6)
LPE(18:1)
PI (pmole/mg protein)
PI(36:4)
PI(38:5)
PI(38:4)
PI(32:1)
PI(40:6)
PI(38:6)
PI(36:1)
PI(34:1)
PI(34:2)
PI(38:3)
PI(36:2)
PI(36:3)
PI(36:4)
PI(38:5)
PI(38:4)
PG (pmole/mg protein)
PG(36:1)
PG(36:3)
PG(36:4)
PG(34:1)
PG(32:0)
PG(38:6)
PG(34:3)
PG(36:1)
PG(36:3)
PG(36:4)
PG(34:1)
PEp (pmole/mg protein)
PE (18:0p/22:6)
PE (16:0p/20:4)
PE (16:0p/22:6)
PE (18:0p/20:4)
PE(18:0p/18:3)
PE(16:0p/20:3)
PE(18:0p/20:3)
PE(16:0p/22:4)
PE(16:0p/18:2)
PE(18:0p/20:5)
PE(18:0p/18:2)
PE(16:0p/22:5)
PE(18:0p/18:1)
PE(18:0p/22:6)
PE(16:0p/20:4)
PE(16:0p/22:6)
PE(18:0p/20:4)

### Figure S2

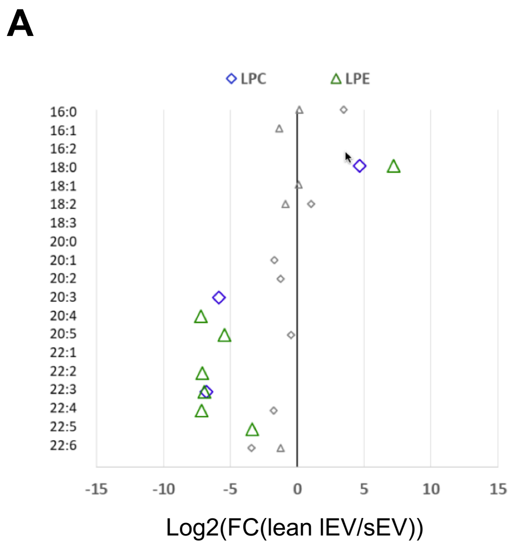
A
LPC
LPE
16:0
16:1
16:2
18:0
18:1
18:2
18:3
20:0
20:1
20:2
20:3
20:4
20:5
22:1
22:2
22:3
22:4
22:5
22:6
-15
-10
-5
0
5
10
15
Log2(FC(lean lEV/sEV))

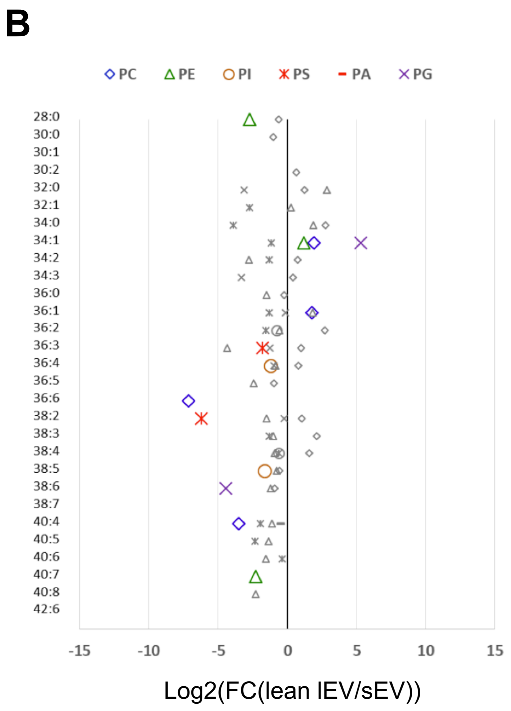
B
PC
PE
PI
PS
PA
PG
28:0
30:0
30:1
30:2
32:0
32:1
34:0
34:1
34:2
34:3
36:0
36:1
36:2
36:3
36:4
36:5
36:6
38:2
38:3
38:4
38:5
38:6
38:7
40:4
40:5
40:6
40:7
40:8
42:6
-15
-10
-5
0
5
10
15
Log2(FC(lean lEV/sEV))

### Figure S3

**A** lEV

**B** sEV

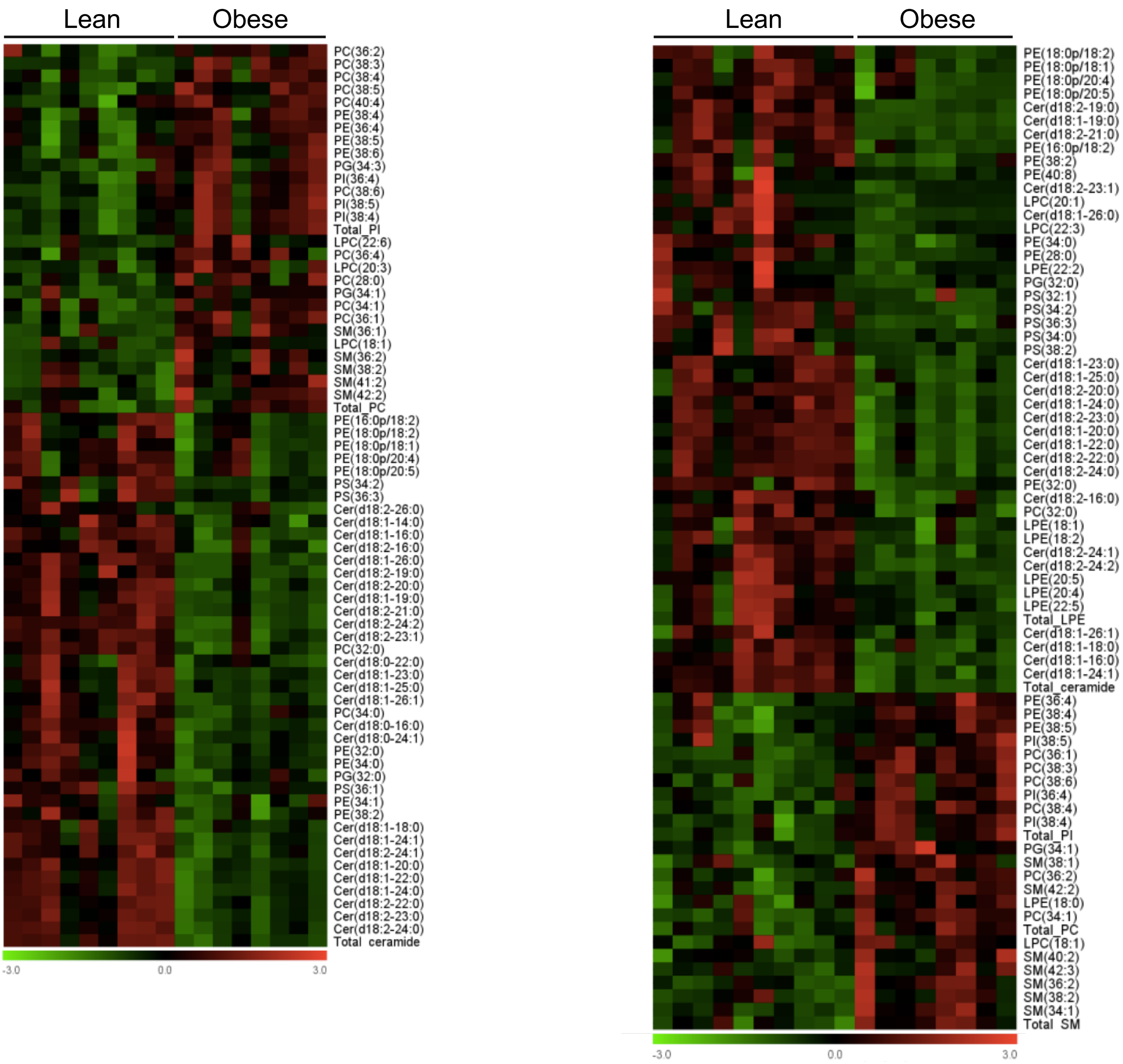

### Figure S4

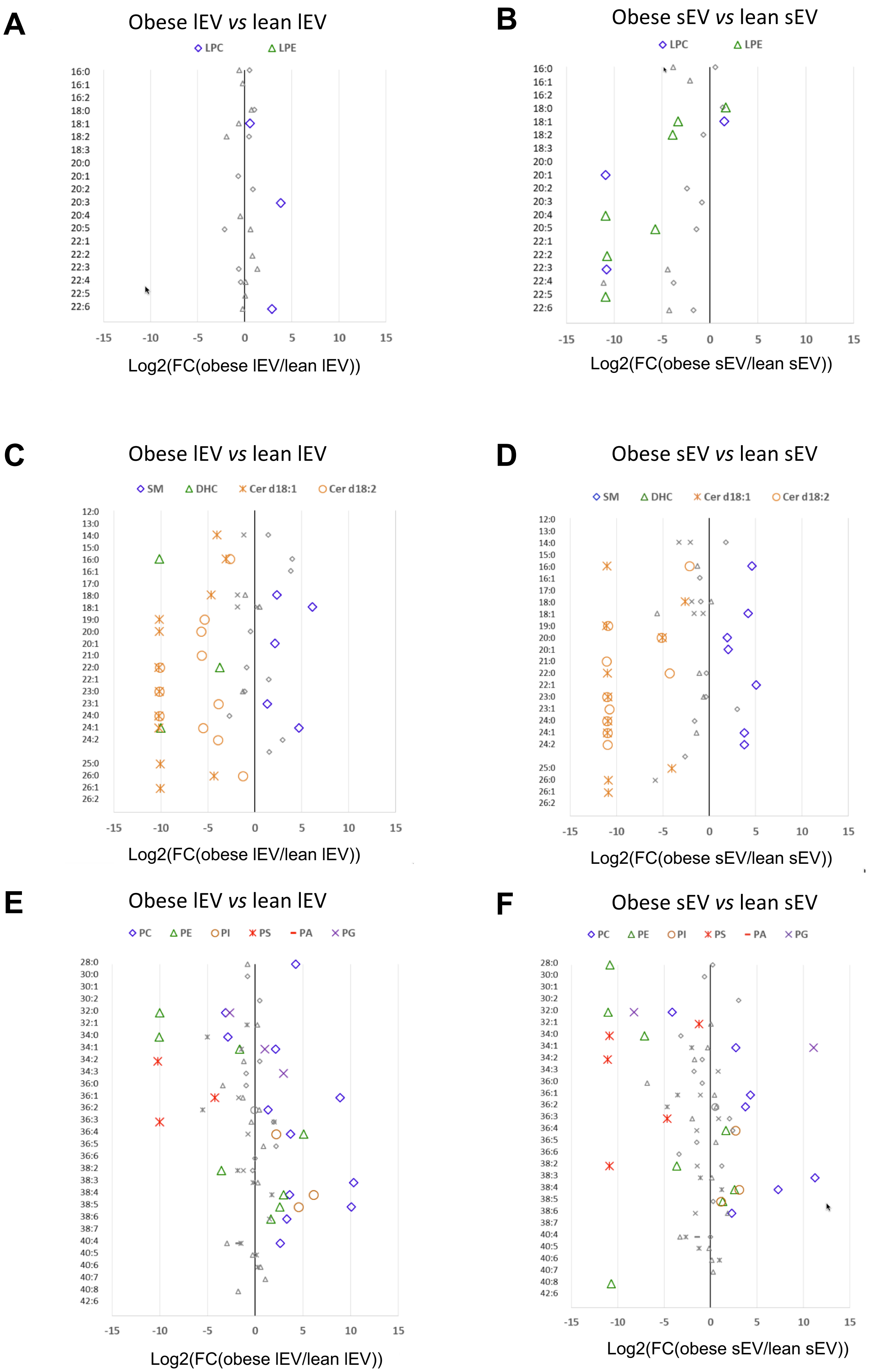
A
Obese lEV vs lean lEV
LPC
LPE
Log2(FC(obese lEV/lean lEV))
B
Obese sEV vs lean sEV
Log2(FC(obese sEV/lean sEV))
C
SM
DHC
Cer d18:1
Cer d18:2
D
E
PC
PE
PI
PS
PA
PG
F

**Table S1 : Phenotypic characteristics of obese and lean mice**

| | Lean | Obese (*ob/ob*) |
|---|---|---|
| Total weight (g) | 27.2 ± 2.1 | 47.1 ± 6.8**** |
| SAT weight (g) | 0.41 ± 0.13 | 4.91 ± 1.68**** |
| VAT weight (g) | 0.80 ± 0.19 | 3.74 ± 0.98*** |
| Muscle weight (g) | 0.44 ± 0.007 | 0.27 ± 0.02*** |
| Liver weight (g) | 1.23 ± 0.15 | 3.67 ± 0.8**** |
| Glycemia (mg/dL) | 135 ± 7 | 202 ± 23* |
| Insulinemia | 0.82 ± 0.28 | 3.54 ± 0.60**** |

### Table S2 : Structural elucidation

**EV, VAT**

| Lipid class | Lipid subclass | Lipid species | MS experiment for strcutural elucidation | sEV | lEV | VAT |
|---|---|---|---|---|---|---|
| | LysoPC | LPC(16:0) | [M+H]+ PIS 184m/z | | | |
| | | LPC(16:1) | | | | |
| | | LPC(16:2) | | | | |
| | | LPC(18:0) | | | | |
| | | LPC(18:1) | | | | |
| | | LPC(18:2) | | | | |
| | | LPC(18:3) | | | | |
| | | LPC(20:0) | | | | |
| | | LPC(20:1) | | | | |
| | | LPC(20:2) | | | | |
| | | LPC(20:3) | | | | |
| | | LPC(20:4) | | | | |
| | | LPC(20:5) | | | | |
| | | LPC(22:3) | | | | |
| | | LPC(22:4) | | | | |
| | | LPC(22:5) | | | | |
| | | LPC(22:6) | | | | |
| | LysoPE | LPE(16:0) | [M+H]+ NL 141 | | | |
| | | LPE(16:1) | | | | |
| | | LPE(18:0) | | | | |
| | | LPE(18:1) | | | | |
| | | LPE(18:2) | | | | |
| | | LPE(18:3) | | | | |
| | | LPE(20:3) | | | | |
| | | LPE(20:4) | | | | |
| | | LPE(20:5) | | | | |
| | | LPE(22:1) | | | | |
| | | LPE(22:2) | | | | |
| | | LPE(22:3) | | | | |
| | | LPE(22:4) | | | | |

**Plasma**

| Lipid class | Lipid subclass | Lipid species | MS experiment for strcutural elucidation | elucidated lipid in plasma |
|---|---|---|---|---|
| | LysoPC | LPC(16:0) | [M+H]+ PIS 184m/z | |
| | | LPC(16:1) | | |
| | | LPC(18:0) | | |
| | | LPC(18:1) | | |
| | | LPC(18:2) | | |
| | | LPC(18:3) | | |
| | | LPC(20:3) | | |
| | | LPC(20:4) | | |
| | | LPC(20:5) | | |
| | | LPC(22:5) | | |
| | | LPC(22:6) | | |
| | LysoPE | LPE(16:0) | [M+H]+ NL 141 | |
| | | LPE(16:1) | | |
| | | LPE(18:0) | | |
| | | LPE(18:1) | | |
| | | LPE(18:2) | | |
| | | LPE(18:3) | | |
| | | LPE(20:3) | | |
| | | LPE(20:4) | | |
| | | LPE(20:5) | | |
| | | LPE(22:1) | | |
| | | LPE(22:4) | | |
| | | LPE(22:5) | | |
| | | LPE(22:6) | | |
| | Phosphatidic acid | PA(34:2) | Extrapolated structural elucidation | 16:0_18:2 |
| | | PA(36:1) | | 18:0_18:1 |
| | | PA(36:2) | | 18:0_18:2 |
| | | PA(36:4) | | 16:0_20:4 |
| | | PC(28:0) | | 14:0_14:0 |
| | | PC(30:0) | | 14:0_16:0 |

| | | | | | | |
|---|---|---|---|---|---|---|
| | | LPE(22:5) | | | | |
| | | LPE(22:6) | | | | |
| | Phosphatidic acid | PA(32:0) | Extrapolated structural elucidation | 16:0_16:0 | 16:0_16:1 | 16:0_16:2 |
| | | PA(36:3) | | 18:1_18:2 | 18:1_18:3 | 18:1_18:4 |
| | | PA(36:4) | | 16:0_20:4 | 16:0_20:5 | 16:0_20:6 |
| | | PA(38:4) | | 18:0_20:4 | 18:0_20:5 | 18:0_20:6 |
| | | PA(38:5) | | 16:0_22:5 | 16:0_22:6 | 16:0_22:7 |
| | | PA(40:4) | | 20:0_20:4 | 20:0_20:5 | 20:0_20:6 |
| | | PA(40:7) | | 18:1_22:6 | 18:1_22:7 | 18:1_22:8 |
| | Phosphatidylcholine | PC(28:0) | Extrapolated structural elucidation | 14:0_14:0 | 14:0_14:1 | 14:0_14:2 |
| | | PC(30:0) | [M-H]- PIS FA chain | 14:0_16:0 | 14:0_16:0 | 14:0_16:0 |
| | | PC(30:1) | Extrapolated structural elucidation | 14:0_16:1 | 14:0_16:2 | 14:0_16:3 |
| | | PC(30:2) | | 14:1_16:2 | 14:1_16:3 | 14:1_16:4 |
| | | PC(32:0) | [M-H]- PIS FA chain | 16:0_16:0 | 16:0_16:0 | 16:0_16:0 |
| | | PC(32:1) | Extrapolated structural elucidation | 16:0_16:1 | 16:0_16:2 | 16:0_16:3 |
| | | PC(34:0) | | 16:0_18:0 | 16:0_18:0 | 16:0_18:0 |
| | | PC(34:1) | | 16:0_18:1 | 16:0_18:1 | 16:0_18:1 |
| | | PC(34:2) | [M-H]- PIS FA chain | 18:2_16:0 | 18:2_16:0 | 18:2_16:0 |
| | | PC(34:3) | | 18:2_16:1 | 18:2_16:1 | 18:2_16:1 |
| | | PC(36:0) | | 18:0_18:0 | 18:0_18:0 | 18:0_18:0 |
| | | PC(36:1) | | 18:1_18:0 | 18:1_18:0 | 18:1_18:0 |
| | | PC(36:2) | | 18:2_18:0 | 18:2_18:0 | 18:2_18:0 |
| | | PC(36:3) | | 18:2_18:1 | 18:2_18:1 | 18:2_18:1 |
| | | PC(36:4) | | 20:4_16:0 | 20:4_16:0 | 20:4_16:0 |
| | | PC(36:5) | | 20:5_16:0 | 20:4_16:1 | 20:5_16:0 |
| | | PC(36:6) | Extrapolated structural elucidation | 14:0_22:6 | 14:0_22:7 | 14:0_22:8 |
| | | PC(38:2) | | 18:0_20:2 | 18:0_20:3 | 18:0_20:4 |
| | | PC(38:3) | [M-H]- PIS FA chain | 20:3_18:0 | 20:3_18:0 | 20:3_18:0 |
| | | PC(38:4) | | 20:4_18:0 | 20:4_18:0 | 20:4_18:0 |
| | | PC(38:5) | | 20:4_18:1 | 20:4_18:1 | 20:4_18:1 |
| | | PC(38:6) | | 22:6_16:0 | 22:6_16:0 | 22:6_16:0 |

| | | | | |
|---|---|---|---|---|
| Glycerophospholipids | Phosphatidylcholine | PC(30:1) | Extrapolated structural elucidation | 14:0_16:1 |
| | | PC(30:2) | | 14:1_16:2 |
| | | PC(32:0) | | 16:0_16:0 |
| | | PC(32:1) | | 16:0_16:1 |
| | | PC(34:1) | | 16:0_18:1 |
| | | PC(34:2) | | 16:0_18:2 |
| | | PC(34:3) | | 16:1_18:2 |
| | | PC(36:0) | | 18:0_18:0 |
| | | PC(36:1) | | 18:0_18:1 |
| | | PC(36:2) | | 18:0_18:2 |
| | | PC(36:3) | | 18:1_18:2 |
| | | PC(36:4) | | 16:0_20:4 |
| | | PC(36:5) | | 16:1_20:4 |
| | | PC(36:6) | | 14:0_22:6 |
| | | PC(38:2) | | 18:0_20:2 |
| | | PC(38:3) | | 18:0_20:3 |
| | | PC(38:4) | | 18:0_20:4 |
| | | PC(38:5) | | 16:0_22:5 |
| | | PC(38:6) | | 16:0_22:6 |
| | | PC(40:4) | | 20:0_20:4 |
| | | PC(40:5) | | 18:0_22:5 |
| | | PC(40:6) | | 18:0_22:6 |
| | | PC(40:7) | | 18:1_22:6 |
| | | PC(40:8) | | 18:2_22:6 |
| | | PE(28:0) | | 14:0_14:0 |
| | | PE(30:0) | | 14:0_16:0 |
| | | PE(30:1) | | 14:0_16:1 |
| | | PE(32:1) | | 16:0_16:1 |
| | | PE(34:1) | | 16:0_18:1 |
| | | PE(34:2) | | 16:0_18:2 |
| | | PE(36:0) | | 18:0_18:0 |

| | | | | | | |
|---|---|---|---|---|---|---|
| Glycerophospholipids | | PC(40:4) | Extrapolated structural elucidation | 20:0_20:4 | 20:0_20:5 | 20:0_20:6 |
| | | PC(40:5) | | 22:5_18:0 | 20:4_20:1 | 22:5_18:0 |
| | | PC(40:6) | [M-H]- PIS FA chain | 22:6_18:0 | 22:6_18:0 | 22:6_18:0 |
| | | PC(40:7) | | 22:6_18:1 | 22:6_18:1 | 22:6_18:1 |
| | | PC(40:8) | | 20:4_20:4 | 20:4_20:4 | 22:6_18:2 |
| | Plasmalogen PE | PE(16:0p/18:1) | [M+H]+ PIS(RCO+PE) | | | |
| | | PE(16:0p/18:2) | | | | |
| | | PE(16:0p/20:1) | | | | |
| | | PE(16:0p/20:2) | | | | |
| | | PE(16:0p/20:3) | | | | |
| | | PE(16:0p/20:4) | | | | |
| | | PE(16:0p/22:4) | | | | |
| | | PE(16:0p/22:5) | | | | |
| | | PE(16:0p/22:6) | | | | |
| | | PE(18:0p/18:1) | | | | |
| | | PE(18:0p/18:2) | | | | |
| | | PE(18:0p/18:3) | | | | |
| | | PE(18:0p/18:4) | | | | |
| | | PE(18:0p/20:3) | | | | |
| | | PE(18:0p/20:4) | | | | |
| | | PE(18:0p/20:5) | | | | |
| | | PE(18:0p/22:6) | | | | |
| | | PE(18:0p/24:5) | | | | |
| | | PE(18:0p/24:6) | | | | |
| | amine | PE(28:0) | Extrapolated structural elucidation | 14:0_14:0 | 14:0_14:1 | 14:0_14:2 |
| | | PE(32:0) | | 16:0_16:0 | 16:0_16:0 | 16:0_16:0 |
| | | PE(32:1) | | 16:0_16:1 | 16:0_16:1 | 16:0_16:1 |
| | | PE(34:0) | | 16:0_18:0 | 16:0_18:1 | 16:0_18:2 |
| | | PE(34:1) | [M-H]- PIS FA chain | 16:0_18:1 | 16:0_18:1 | 16:0_18:1 |
| | | PE(34:2) | | 18:2_16:0 | 18:2_16:0 | 18:2_16:0 |
| | | PE(36:0) | Extrapolated structural elucidation | 18:0_18:0 | 18:0_18:1 | 18:0_18:2 |
| | | PE(36:1) | | 18:1_18:0 | 18:1_18:0 | 18:1_18:0 |
| | | PE(36:2) | [M-H]- PIS FA chain | 18:2_18:0 | 18:2_18:0 | 18:2_18:0 |
| | | PE(36:3) | | 18:2_18:1 | 18:2_18:1 | 18:2_18:1 |

| | | | |
|---|---|---|---|
| Phosphatidylethanolamine | PE(36:1) | Extrapolated structural elucidation | 18:0_18:1 |
| | PE(36:2) | | 18:0_18:2 |
| | PE(36:3) | | 18:1_18:2 |
| | PE(36:4) | | 16:0_20:4 |
| | PE(36:5) | | 16:1_20:4 |
| | PE(38:2) | | 18:0_20:2 |
| | PE(38:3) | | 18:0_20:3 |
| | PE(38:4) | | 18:0_20:4 |
| | PE(38:5) | | 16:0_22:5 |
| | PE(38:6) | | 16:0_22:6 |
| | PE(40:4) | | 20:0_20:4 |
| | PE(40:6) | | 18:0_22:6 |
| | PE(40:7) | | 18:1_22:6 |
| | PE(40:8) | | 18:2_22:6 |
| plasmalogen PE | PE(16:0p/18:2) | [M+H]+ PIS(RCO+PE) | |
| | PE(16:0p/20:1) | | |
| | PE(16:0p/20:2) | | |
| | PE(16:0p/20:3) | | |
| | PE(16:0p/20:4) | | |
| | PE(16:0p/22:4) | | |
| | PE(16:0p/22:5) | | |
| | PE(16:0p/22:6) | | |
| | PE(18:0p/18:1) | | |
| | PE(18:0p/18:2) | | |
| | PE(18:0p/18:3) | | |
| | PE(18:0p/20:3) | | |
| | PE(18:0p/20:4) | | |
| | PE(18:0p/20:5) | | |
| | PE(18:0p/22:6) | | |
| Phosphatidylglycerol | PG(34:1) | Extrapolated structural elucidation | 16:0_18:1 |
| | PG(36:1) | | 18:0_18:1 |
| | PG(36:2) | | 18:0_18:2 |
| | PI(32:0) | | 16:0_16:0 |
| | PI(32:1) | | 16:0_16:1 |

| | | | | | | |
|---|---|---|---|---|---|---|
| | Phosphatidylethanol | PE(36:4) | chain | 20:4_16:0 | 20:4_16:0 | 20:4_16:0 |
| | | PE(36:5) | | 18:3_18:2 | 18:3_18:2 | 18:3_18:2 |
| | | PE(38:2) | Extrapolated structural elucidation | 18:0_20:2 | 18:0_20:3 | 18:0_20:4 |
| | | PE(38:3) | [M-H]- PIS FA chain | 20:3_18:0 | 20:3_18:0 | 20:3_18:0 |
| | | PE(38:4) | Extrapolated structural elucidation | 18:0_20:4 | 18:0_20:5 | 18:0_20:6 |
| | | PE(38:5) | [M-H]- PIS FA chain | 20:4_18:1 | 20:4_18:1 | 20:4_18:1 |
| | | PE(38:6) | | 22:6_16:0 | 22:6_16:0 | 22:6_16:0 |
| | | PE(40:4) | Extrapolated structural elucidation | 20:0_20:4 | 20:0_20:5 | 20:0_20:6 |
| | | PE(40:5) | | 18:0_22:5 | 18:0_22:5 | 18:0_22:5 |
| | | PE(40:6) | | 18:0_22:6 | 18:0_22:6 | 18:0_22:6 |
| | | PE(40:7) | | 18:1_22:6 | 18:1_22:7 | 18:1_22:8 |
| | | PE(40:8) | | 18:2_22:6 | 18:2_22:7 | 18:2_22:8 |
| | Phosphatidylglycerol | PG(32:0) | Extrapolated structural elucidation | 16:0_16:0 | 16:0_16:1 | 16:0_16:2 |
| | | PG(34:1) | | 16:0_18:1 | 16:0_18:2 | 16:0_18:3 |
| | | PG(34:3) | | 16:1_18:2 | 16:1_18:3 | 16:1_18:4 |
| | | PG(36:1) | | 18:0_18:1 | 18:0_18:2 | 18:0_18:3 |
| | | PG(36:3) | | 18:1_18:2 | 18:1_18:3 | 18:1_18:4 |
| | | PG(36:4) | | 16:0_20:4 | 16:0_20:5 | 16:0_20:6 |
| | | PG(38:2) | | 18:0_20:2 | 18:0_20:2 | 18:0_20:2 |
| | | PG(38:3) | | 18:0_20:3 | 18:0_20:3 | 18:0_20:3 |
| | | PG(38:6) | | 16:0_22:6 | 16:0_22:7 | 16:0_22:8 |
| | Phosphatidylinositol | PI(32:1) | Extrapolated structural elucidation | 16:0_16:1 | 16:0_16:2 | 16:0_16:3 |
| | | PI(34:1) | | 16:0_18:1 | 16:0_18:1 | 16:0_18:1 |
| | | PI(34:2) | | 16:0_18:2 | 16:0_18:2 | 16:0_18:2 |
| | | PI(36:1) | | 18:0_18:1 | 18:0_18:2 | 18:0_18:3 |
| | | PI(36:2) | [M-H]- PIS FA chain | 18:1_18:1 | 18:1_18:1 | 18:1_18:1 |
| | | PI(36:3) | | 18:2_18:1 | 18:2_18:1 | 18:2_18:1 |
| | | PI(36:4) | | 20:4_16:0 | 20:4_16:0 | 20:4_16:0 |
| | | PI(38:3) | | 20:3_18:0 | 20:3_18:0 | 20:3_18:0 |
| | | PI(38:4) | | 20:4_18:0 | 20:4_18:0 | 20:4_18:0 |
| | | PI(38:5) | | 20:5_18:0 | 20:5_18:0 | 20:5_18:0 |
| | | PI(38:6) | | 22:6_16:0 | 22:6_16:0 | 22:6_16:0 |

| | | | | |
|---|---|---|---|---|
| | Phosphatidylinositol | PI(34:1) | Extrapolated structural elucidation | 16:0_18:1 |
| | | PI(36:1) | | 18:0_18:1 |
| | | PI(36:2) | | 18:0_18:2 |
| | | PI(36:3) | | 18:1_18:2 |
| | | PI(36:4) | | 16:0_20:4 |
| | | PI(38:3) | | 18:0_20:3 |
| | | PI(38:4) | | 18:0_20:4 |
| | | PI(38:5) | | 16:0_22:5 |
| | | PI(38:6) | | 16:0_22:6 |
| | | PI(40:4) | | 20:0_20:4 |
| | | PI(40:5) | | 18:0_22:5 |
| | | PI(40:6) | | 18:0_22:6 |
| | | PI(40:7) | | 18:1_22:6 |
| | Phosphatidylserine | PS(36:1) | Extrapolated structural elucidation | 18:0_18:1 |
| | | PS(38:4) | | 18:0_20:4 |
| | | PS(40:5) | | 18:0_22:5 |
| | | PS(40:6) | | 18:0_22:6 |
| | Dihydroceramides | Cer(d18:0-16:0) | [M+H]+ PIS 266m/z | |
| | | Cer(d18:0-18:0) | | |
| | | Cer(d18:0-18:1) | | |
| | | Cer(d18:0-19:0) | | |
| | | Cer(d18:0-20:0) | | |
| | | Cer(d18:0-24:0) | | |
| | | Cer(d18:0-24:1) | | |
| | | Cer(d18:0-25:0) | | |
| | | Cer(d18:0-26:0) | | |
| | | Cer(d18:0-26:1) | | |
| | | Cer(d18:0-26:2) | | |
| | | Cer(d18:1-14:0) | | |
| | | Cer(d18:1-16:0) | | |
| | | Cer(d18:1-18:0) | | |
| | | Cer(d18:1-18:1) | | |

| | | | | | | |
|---|---|---|---|---|---|---|
| | | PI(40:6) | Extrapolated structural elucidation | 18:0_22:6 | 18:0_22:7 | 18:0_22:8 |
| | Phosphatidylserine | PS(32:0) | Extrapolated structural elucidation | 16:0_16:0 | 16:0_16:0 | 16:0_16:0 |
| | | PS(32:1) | | 16:0_16:1 | 16:0_16:1 | 16:0_16:1 |
| | | PS(34:0) | | 16:0_18:0 | 16:0_18:0 | 16:0_18:0 |
| | | PS(34:1) | | 16:0_18:1 | 16:0_18:1 | 16:0_18:1 |
| | | PS(34:2) | | 16:0_18:2 | 16:0_18:2 | 16:0_18:2 |
| | | PS(36:1) | | 18:0_18:1 | 18:0_18:1 | 18:0_18:1 |
| | | PS(36:2) | | 18:0_18:2 | 18:0_18:2 | 18:0_18:2 |
| | | PS(36:3) | | 18:1_18:2 | 18:1_18:3 | 18:1_18:4 |
| | | PS(36:5) | | 16:1_20:4 | 16:1_20:5 | 16:1_20:6 |
| | | PS(38:2) | | 18:0_20:2 | 18:0_20:2 | 18:0_20:2 |
| | | PS(38:3) | | 18:0_20:3 | 18:0_20:3 | 18:0_20:3 |
| | | PS(38:4) | | 18:0_20:4 | 18:0_20:4 | 18:0_20:4 |
| | | PS(40:4) | | 20:0_20:4 | 20:0_20:5 | 20:0_20:6 |
| | | PS(40:5) | | 18:0_22:5 | 18:0_22:5 | 18:0_22:5 |
| | | PS(40:6) | | 18:0_22:6 | 18:0_22:6 | 18:0_22:6 |
| | Dihydroceramides (DHC) | Cer(d18:0/13:0) | [M+H]+ PIS 266m/z | | | |
| | | Cer(d18:0/14:0) | | | | |
| | | Cer(d18:0/15:0) | | | | |
| | | Cer(d18:0/16:0) | | | | |
| | | Cer(d18:0/17:0) | | | | |
| | | Cer(d18:0/18:0) | | | | |
| | | Cer(d18:0/18:1) | | | | |
| | | Cer(d18:0/19:0) | | | | |
| | | Cer(d18:0/20:0) | | | | |
| | | Cer(d18:0/22:0) | | | | |
| | | Cer(d18:0/23:0) | | | | |
| | | Cer(d18:0/24:0) | | | | |
| | | Cer(d18:0/24:1) | | | | |
| | | Cer(d18:0/25:0) | | | | |
| | | Cer(d18:0/26:0) | | | | |
| | | Cer(d18:0/26:1) | | | | |
| | | Cer(d18:0/26:2) | | | | |
| | | Cer(d18:1/12:0) | | | | |
| | | Cer(d18:1/13:0) | | | | |
| | | Cer(d18:1/14:0) | | | | |

| | | | | |
|---|---|---|---|---|
| sphingolipids | Ceramides | Cer(d18:1-19:0) | [M+H]+ PIS 264m/z | |
| | | Cer(d18:1-20:0) | | |
| | | Cer(d18:1-22:0) | | |
| | | Cer(d18:1-23:0) | | |
| | | Cer(d18:1-24:0) | | |
| | | Cer(d18:1-24:1) | | |
| | | Cer(d18:1-25:0) | | |
| | | Cer(d18:1-26:0) | | |
| | | Cer(d18:1-26:1) | | |
| | Sphingadienine ceramides | Cer(d18:2-14:0) | [M+H]+ PIS 262m/z | |
| | | Cer(d18:2-16:0) | | |
| | | Cer(d18:2-18:0) | | |
| | | Cer(d18:2-18:1) | | |
| | | Cer(d18:2-19:0) | | |
| | | Cer(d18:2-20:0) | | |
| | | Cer(d18:2-21:0) | | |
| | | Cer(d18:2-22:0) | | |
| | | Cer(d18:2-23:0) | | |
| | | Cer(d18:2-23:1) | | |
| | | Cer(d18:2-24:0) | | |
| | | Cer(d18:2-24:1) | | |
| | | Cer(d18:2-24:2) | | |
| | | Cer(d18:2-26:0) | | |
| | | Cer(d18:2-26:1) | | |
| | Sphingomyelin | SM(30:1) | Extrapolated structural elucidation | 14:0 |
| | | SM(32:1) | | 16:0 |
| | | SM(34:1) | | 16:0 |
| | | SM(34:2) | | 16:1 |
| | | SM(36:1) | | 18:0 |
| | | SM(36:2) | | 18:0 |
| | | SM(38:1) | | 20:0 |
| | | SM(38:2) | | 20:0 |
| | | SM(40:1) | | 22:0 |
| | | SM(40:2) | | 22:0 |
| | | SM(41:1) | | 23:0 |
| | | SM(41:2) | | 23:0 |

| | | | | | | |
|---|---|---|---|---|---|---|
| sphingolipids | Ceramides | Cer(d18:1/15:0) | [M+H]+ PIS 264m/z | | | |
| | | Cer(d18:1/16:0) | | | | |
| | | Cer(d18:1/18:0) | | | | |
| | | Cer(d18:1/18:1) | | | | |
| | | Cer(d18:1/19:0) | | | | |
| | | Cer(d18:1/20:0) | | | | |
| | | Cer(d18:1/22:0) | | | | |
| | | Cer(d18:1/23:0) | | | | |
| | | Cer(d18:1/24:0) | | | | |
| | | Cer(d18:1/24:1) | | | | |
| | | Cer(d18:1/25:0) | | | | |
| | | Cer(d18:1/26:0) | | | | |
| | | Cer(d18:1/26:1) | | | | |
| | Sphingadienine ceramides | Cer(d18:2/13:0) | [M+H]+ PIS 262m/z | | | |
| | | Cer(d18:2/14:0) | | | | |
| | | Cer(d18:2/15:0) | | | | |
| | | Cer(d18:2/16:0) | | | | |
| | | Cer(d18:2/18:0) | | | | |
| | | Cer(d18:2/18:1) | | | | |
| | | Cer(d18:2/19:0) | | | | |
| | | Cer(d18:2/20:0) | | | | |
| | | Cer(d18:2/20:1) | | | | |
| | | Cer(d18:2/21:0) | | | | |
| | | Cer(d18:2/22:0) | | | | |
| | | Cer(d18:2/23:0) | | | | |
| | | Cer(d18:2/23:1) | | | | |
| | | Cer(d18:2/24:0) | | | | |
| | | Cer(d18:2/24:1) | | | | |
| | | Cer(d18:2/24:2) | | | | |
| | | Cer(d18:2/26:0) | | | | |
| | | Cer(d18:2/26:1) | | | | |
| | yelin | SM(30:1) | Extrapolated | C14:0 | C14:1 | C14:2 |
| | | SM(32:1) | | C16:0 | C16:0 | C16:0 |
| | | SM(34:1) | | C16:0 | C16:0 | C16:0 |
| | | SM(34:2) | | C16:1 | C16:1 | C16:1 |
| | | SM(36:1) | | C18:0 | C18:0 | C18:0 |
| | | SM(36:2) | | C18:0 | C18:0 | C18:0 |
| | | SM(38:1) | | C20:0 | C20:0 | C20:0 |

| | | | | |
|---|---|---|---|---|
| | | SM(42:1) | | 24:0 |
| | | SM(42:2) | | 24:1 |
| | | SM(42:3) | | 24:1 |
| | | SM(42:4) | | 24:2 |
| | Cholesteryl esters | CE(16:0) | [M+NH4]+ PIS 369m/z | 16:0 |
| | | CE(16:1) | | 16:1 |
| | | CE(18:0) | | 18:0 |
| | | CE(18:1) | | 18:1 |
| | | CE(18:2) | | 18:2 |
| | | CE(18:3) | | 18:3 |
| | | CE(20:3) | | 20:3 |
| | | CE(20:4) | | 20:4 |
| | | CE(22:5) | | 22:5 |
| | | CE(22:6) | | 22:6 |
| | Diacylglycerols | DG(14:0_16:0) | [M+NH4]+ NL(RCOO+NH3) | 14:0 |
| | | DG(14:0_16:1) | | 14:0 |
| | | DG(16:0_16:0) | | 16:0 |
| | | DG(14:0_18:1) | | 14:0 |
| | | DG(14:0_18:2) | | 14:0 |
| | | DG(16:0_18:0) | | 16:0 |
| | | DG(16:0_18:1) | | 16:0 |
| | | DG(16:0_18:2) | | 16:0 |
| | | DG(16:1_18:1) | | 16:1 |
| | | DG(16:1_18:2) | | 16:1 |
| | | DG(16:0_18:3) | | 16:0 |
| | | DG(16:1_18:3) | | 16:1 |
| | | DG(18:0_18:0) | | 18:0 |
| | | DG(18:0_18:1) | | 18:0 |
| | | DG(18:0_18:2) | | 18:0 |
| | | DG(18:1_18:1) | | 18:1 |
| | | DG(16:0_20:3) | | 16:0 |
| | | DG(16:0_20:4) | | 16:0 |
| | | DG(18:1_18:3) | | 18:1 |
| | | DG(18:2_18:2) | | 18:2 |
| | | DG(16:1_20:4) | | 16:1 |
| | | DG(18:2_18:3) | | 18:2 |
| | | DG(16:0_20:5) | | 16:0 |
| | | DG(18:1_20:3) | | 18:1 |

| | | | | | | |
|---|---|---|---|---|---|---|
| | Sphingomy | SM(38:2) | Extrapolated structural elucidation | C20:0 | C20:0 | C20:0 |
| | | SM(40:1) | | C22:0 | C22:0 | C22:0 |
| | | SM(40:2) | | C22:0 | C22:0 | C22:0 |
| | | SM(41:1) | | C23:0 | C23:0 | C23:0 |
| | | SM(41:2) | | C23:0 | C23:0 | C23:0 |
| | | SM(42:1) | | C24:0 | C24:0 | C24:0 |
| | | SM(42:2) | | C24:1 | C24:1 | C24:1 |
| | | SM(42:3) | | C24:1 | C24:1 | C24:1 |
| | | SM(42:4) | | C24:2 | C24:2 | C24:2 |
| | Cholesteryl esters | CE(16:0) | [M+NH4]+ PIS 369m/z | | | |
| | | CE(16:1) | | | | |
| | | CE(18:0) | | | | |
| | | CE(18:1) | | | | |
| | | CE(18:2) | | | | |
| | | CE(18:3) | | | | |
| | | CE(20:3) | | | | |
| | | CE(20:4) | | | | |
| | | CE(22:5) | | | | |
| | | CE(22:6) | | | | |
| | acylglycerols | DG(14:0_16:0) | [M+NH4]+ NL(RCOO+NH3) | | | |
| | | DG(14:0_16:1) | | | | |
| | | DG(14:0_18:1) | | | | |
| | | DG(14:0_18:2) | | | | |
| | | DG(16:0_16:0) | | | | |
| | | DG(16:0_18:0) | | | | |
| | | DG(16:0_18:1) | | | | |
| | | DG(16:0_18:2) | | | | |
| | | DG(16:0_18:3) | | | | |
| | | DG(16:0_18:4) | | | | |
| | | DG(16:0_20:3) | | | | |
| | | DG(16:0_20:4) | | | | |
| | | DG(16:0_20:5) | | | | |
| | | DG(16:0_22:5) | | | | |
| | | DG(16:0_22:6) | | | | |
| | | DG(16:1_18:1) | | | | |
| | | DG(16:1_18:2) | | | | |
| | | DG(16:1_18:3) | | | | |
| | | DG(16:1_20:4) | | | | |

| | | | | |
|---|---|---|---|---|
| Neutral lipids | | DG(16:0_22:5) | | 16:0 |
| | | DG(18:1_20:4) | | 18:1 |
| | | DG(16:0_22:6) | | 16:0 |
| | | DG(18:2_20:4) | | 18:2 |
| | | DG(18:1_22:6) | | 18:1 |
| | Triacylglycerols | TG(48:0-16:0/32:0 | [M+NH4]+ NL(RCOO+NH3) + extrapolated 2nd and 3rd chains | TG(16:0_16:0_16:0) |
| | | TG(48:1-16:1/32:0 | | TG(16:0_16:0_16:1) |
| | | TG(48:1-18:1/30:0 | | TG(14:0_16:0_18:1) |
| | | TG(48:2-14:1/34:1 | | TG(14:1_16:0_18:1) |
| | | TG(48:2-16:0/32:2 | | TG(16:0_16:1_16:1) |
| | | TG(48:2-18:1/30:1 | | TG(14:0_16:1_18:1) |
| | | TG(48:2-18:2/30:0 | | TG(14:0_16:0_18:2) |
| | | TG(48:3-14:0/34:3 | | TG(14:0_16:1_18:2) |
| | | TG(48:3-16:1/32:2 | | TG(16:1_16:1_16:1) |
| | | TG(49:1-14:0/35:1 | | TG(14:0_17:0_18:1) |
| | | TG(49:1-15:0/34:1 | | TG(15:0_16:0_18:1) |
| | | TG(49:1-17:0/32:1 | | TG(16:0_16:1_17:0) |
| | | TG(50:0-18:0/32:0 | | TG(16:0_16:0_18:0) |
| | | TG(50:1-14:0/36:1 | | TG(14:0_18:0_18:1) |
| | | TG(50:1-18:1/32:0 | | TG(16:0_16:0_18:1) |
| | | TG(50:2-18:0/32:2 | | TG(16:1_16:1_18:0) |
| | | TG(50:2-18:1/32:1 | | TG(16:0_16:1_18:1) |
| | | TG(50:2-18:2/32:0 | | TG(16:0_16:0_18:2) |
| | | TG(50:3-14:1/36:2 | | TG(14:1_18:1_18:1) |
| | | TG(50:3-18:1/32:2 | | TG(16:1_16:1_18:1) |
| | | TG(50:3-18:2/32:1 | | TG(16:1_16:0_18:2) |
| | | TG(50:4-14:0/36:4 | | TG(14:0_16:0_20:4) |
| | | TG(50:4-18:2/32:2 | | TG(16:1_16:1_18:2) |
| | | TG(51:1-18:1/33:0 | | TG(16:0_17:0_18:1) |
| | | TG(51:2-15:0/36:2 | | TG(15:0_18:1_18:1) |
| | | TG(51:2-16:0/35:2 | | TG(16:0_17:0_18:2) |
| | | TG(52:1-18:1/34:0 | | TG(16:0_18:0_18:1) |
| | | TG(52:2-16:0/36:2 | | TG(16:0_18:1_18:1) |
| | | TG(52:3-16:1/36:2 | | TG(16:1_18:1_18:1) |
| | | TG(52:3-18:2/34:1 | | TG(16:0_18:1_18:2) |
| | | TG(52:4-16:0/36:4 | | TG(16:0_16:0_20:4) |
| | | TG(52:4-16:1/36:3 | | TG(16:1_16:0_20:3) |
| | | TG(52:4-20:4/32:0 | | TG(16:0_16:0_20:4) |

| | | |
|---|---|---|
| neutral lipids | Di | DG(18:0_18:0) |
| | | DG(18:0_18:1) |
| | | DG(18:0_18:2) |
| | | DG(18:0_20:3) |
| | | DG(18:0_20:4) |
| | | DG(18:0_22:6) |
| | | DG(18:1_18:1) |
| | | DG(18:1_18:2) |
| | | DG(18:1_18:3) |
| | | DG(18:1_20:2) |
| | | DG(18:1_20:3) |
| | | DG(18:1_20:4) |
| | | DG(18:1_22:6) |
| | | DG(18:2_18:2) |
| | | DG(18:2_18:3) |
| | | DG(18:2_20:4) |
| | | TG(48:0-16:0_32:0) |
| | | TG(48:1-16:1_32:0) |
| | | TG(48:1-18:1_30:0) |
| | | TG(48:2-14:1_34:1) |
| | | TG(48:2-16:0_32:2) |
| | | TG(48:2-18:1_30:1) |
| | | TG(48:2-18:2_30:0) |
| | | TG(48:3-14:0_34:3) |
| | | TG(48:3-16:1_32:2) |
| | | TG(49:1-14:0_35:1) |
| | | TG(49:1-15:0_34:1) |
| | | TG(49:1-17:0_32:1) |
| | | TG(50:0-18:0_32:0) |
| | | TG(50:1-14:0_36:1) |
| | | TG(50:1-18:1_32:0) |
| | | TG(50:2-18:0_32:2) |
| | | TG(50:2-18:1_32:1) |
| | | TG(50:2-18:2_32:0) |
| | | TG(50:3-14:1_36:2) |
| | | TG(50:3-18:1_32:2) |
| | | TG(50:3-18:2_32:1) |
| | | TG(50:4-14:0_36:4) |

| | | | | |
|---|---|---|---|---|
| | | TG(52:5-20:4/32:1 | | TG(16:0_16:1_20:4) |
| | | TG(53:2-17:0/36:2 | | TG(17:0_18:1_18:1) |
| | | TG(54:1-18:1/36:0 | | TG(18:0_18:0_18:1) |
| | | TG(54:2-18:0/36:2 | | TG(18:0_18:1_18:1) |
| | | TG(54:3-18:1/36:2 | | TG(18:1_18:1_18:1) |
| | | TG(54:4-18:0/36:4 | | TG(18:0_16:0_20:4) |
| | | TG(54:4-18:2/36:2 | | TG(18:1_18:1_18:2) |
| | | TG(54:4-20:4/34:0 | | TG(16:0_18:0_20:4) |
| | | TG(54:5-18:1/36:4 | | TG(18:1_16:0_20:4) |
| | | TG(54:6-18:2/36:4 | | TG(18:2_16:0_20:4) |
| | | TG(54:6-18:3/36:3 | | TG(18:3_16:0_20:3) |
| | | TG(54:6-20:4/34:2 | | TG(16:0_18:2_20:4) |
| | | TG(54:6-22:6/32:0 | | TG(16:0_16:0_22:6) |
| | | TG(56:6-20:4/36:2 | | TG(18:1_18:1_20:4) |
| | | TG(56:6-20:5/36:1 | | TG(18:0_18:1_20:5) |
| | | TG(56:6-22:5/34:1 | | TG(16:0_18:1_22:5) |
| | | TG(56:6-22:6/34:0 | | TG(16:0_18:0_22:6) |
| | | TG(56:8-20:4/36:4 | | TG(16:0_20:4_20:4) |
| | | TG(58:8-22:6/36:2 | | TG(18:1_18:1_22:6) |

| | | | |
|---|---|---|---|
| | Triacylglycerols | TG(50:4-18:2_32:2) | [M+NH4]+ NL(RCOO+NH3) |
| | | TG(51:0-18:0_33:0) | |
| | | TG(51:1-18:1_33:0) | |
| | | TG(51:2-15:0_36:2) | |
| | | TG(51:2-16:0_35:2) | |
| | | TG(51:2-16:1_35:1) | |
| | | TG(52:1-18:1_34:0) | |
| | | TG(52:2-16:0_36:2) | |
| | | TG(52:3-16:1_36:2) | |
| | | TG(52:3-18:2_34:1) | |
| | | TG(52:4-16:0_36:4) | |
| | | TG(52:4-16:1_36:3) | |
| | | TG(52:4-20:4_32:0) | |
| | | TG(52:5-20:4_32:1) | |
| | | TG(53:2-17:0_36:2) | |
| | | TG(54:1-18:1_36:0) | |
| | | TG(54:2-18:0_36:2) | |
| | | TG(54:2-20:2_34:0) | |
| | | TG(54:3-18:1_36:2) | |
| | | TG(54:3-20:3_34:0) | |
| | | TG(54:4-18:0_36:4) | |
| | | TG(54:4-18:2_36:2) | |
| | | TG(54:4-20:4_34:0) | |
| | | TG(54:5-18:1_36:4) | |
| | | TG(54:5-20:4_34:1) | |
| | | TG(54:5-22:5_32:0) | |
| | | TG(54:6-18:2_36:4) | |
| | | TG(54:6-18:3_36:3) | |
| | | TG(54:6-20:4_34:2) | |
| | | TG(54:6-22:6_32:0) | |
| | | TG(56:6-20:4_36:2) | |
| | | TG(56:6-20:5_36:1) | |
| | | TG(56:6-22:5_34:1) | |
| | | TG(56:6-22:6_34:0) | |
| | | TG(56:8-20:4_36:4) | |
| | | TG(58:8-22:6_36:2) | |

**Table S3 : Plasma lipid fingerprints of lean and obese mice**

| | | obese *vs* lean | |
|---|---|---|---|
| | | Log2FC | *p* value |
| **Neutral Lipids** | | | |
| **Cholesterol esters** | CE(16:1) | 0.92818 | 0.0036 |
| | CE(18:1) | 1.106 | 0.0245 |
| | CE(18:3) | 0.27607 | 0.0088 |
| | CE(20:3) | 1.6667 | 0.0071 |
| | CE(20:4) | 1.4249 | 0.0088 |
| | CE(22:5) | 1.6288 | 0.0058 |
| | CE(22:6) | 1.1984 | 0.0054 |
| **Diacylglycerols** | DG(16:0_18:2) | -1.7555 | 0.0118 |
| | DG(16:1_18:1) | -1.2459 | 0.0163 |
| | DG(16:1_18:2) | -3.2715 | 0.0012 |
| | DG(18:2_18:2) | -2.5228 | 0.0049 |
| | DG(18:1_20:3) | 1.2761 | 0.0142 |
| | DG(16:0_22:6) | -1.0062 | 0.0492 |
| | DG(18:2_20:4) | -2.5898 | 0.0116 |
| | DG(18:1_22:6) | -1.8364 | 0.0121 |
| **Triacylglycerols** | TG(48:0-16:0_32:0) | -1.1351 | 0.0488 |
| | TG(48:1-16:1_32:0) | -1.4872 | 0.0362 |
| | TG(48:2-14:1_34:1) | -1.1552 | 0.0379 |
| | TG(48:2-16:0_32:2) | -2.2711 | 0.0033 |
| | TG(48:2-18:1_30:1) | -1.4775 | 0.0154 |
| | TG(48:2-18:2_30:0) | -2.9261 | 0.0030 |
| | TG(48:3-14:0_34:3) | -3.5451 | 0.0045 |
| | TG(48:3-16:1_32:2) | -2.4038 | 0.0043 |
| | TG(49:1-14:0_35:1) | -1.1129 | 0.0040 |
| | TG(49:1-15:0_34:1) | -1.7034 | 0.0140 |
| | TG(50:2-18:0_32:2) | -2.1336 | 0.0025 |
| | TG(50:2-18:1_32:1) | -1.1721 | 0.0143 |
| | TG(50:2-18:2_32:0) | -2.6086 | 0.0040 |
| | TG(50:3-18:1_32:2) | -1.8793 | 0.0132 |
| | TG(50:3-18:2_32:1) | -3.599 | 0.0006 |
| | TG(50:4-14:0_36:4) | -3.5617 | 0.0015 |
| | TG(50:4-18:2_32:2) | -3.522 | 0.0016 |
| | TG(51:1-18:1_33:0) | -1.4596 | 0.0056 |
| | TG(51:2-15:0_36:2) | -1.9691 | 0.0183 |
| | TG(51:2-16:0_35:2) | -1.8693 | 0.0049 |
| | TG(52:3-18:2_34:1) | -2.7666 | 0.0009 |
| | TG(52:4-16:0_36:4) | -3.4432 | 0.0008 |
| | TG(52:4-16:1_36:3) | -2.9304 | 0.0057 |
| | TG(52:4-20:4_32:0) | -0.99352 | 0.0347 |
| | TG(52:5-20:4_32:1) | -2.0387 | 0.0064 |
| | TG(54:1-18:1_36:0) | -2.2306 | 0.0344 |
| | TG(54:4-18:0_36:4) | -2.6158 | 0.0043 |
| | TG(54:4-18:2_36:2) | -2.2552 | 0.0042 |
| | TG(54:5-18:1_36:4) | -2.839 | 0.0020 |
| | TG(54:6-18:2_36:4) | -2.8723 | 0.0065 |
| | TG(54:6-18:3_36:3) | -3.0976 | 0.0024 |
| | TG(54:6-20:4_34:2) | -1.9806 | 0.0038 |
| | TG(54:6-22:6_32:0) | -2.2953 | 0.0478 |
| | TG(56:6-20:5_36:1) | -1.9816 | 0.0018 |
| | TG(56:6-22:5_34:1) | -1.4341 | 0.0105 |
| | TG(56:8-20:4_36:4) | -2.1677 | 0.0021 |
| | TG(58:8-22:6_36:2) | -1.7758 | 0.0159 |
| **Sphingolipids** | | | |
| **Ceramides** | Cer(d18:0/18:0) | 0.86002 | 0.0499 |
| | Cer(d18:0/18:1) | 0.83728 | 0.0468 |
| | Cer(d18:0/20:0) | 0.94519 | 0.0342 |
| | Cer(d18:0/24:1) | 1.4545 | 0.0284 |
| | Cer(d18:1/14:0) | 2.48 | 0.0217 |
| | Cer(d18:1/16:0) | 1.739 | 0.0154 |
| | Cer(d18:1/18:0) | 4.2423 | 0.0273 |
| | Cer(d18:1/18:1) | 2.0894 | 0.0198 |
| | Cer(d18:1/19:0) | 1.3885 | 0.0223 |
| | Cer(d18:1/20:0) | 2.2942 | 0.0007 |
| | Cer(d18:1/23:0) | 0.45342 | 0.0434 |
| | Cer(d18:1/24:1) | 1.475 | 0.0060 |

| | | | |
|---|---|---|---|
| | Cer(d18:1/26:0) | 1.1633 | 0.0211 |
| | Cer(d18:1/26:1) | 1.1369 | 0.0148 |
| | Cer(d18:2/16:0) | 0.96748 | 0.0121 |
| | Cer(d18:2/18:0) | 2.5047 | 0.0135 |
| | Cer(d18:2/18:1) | 1.0745 | 0.0097 |
| | Cer(d18:2/20:1) | 1.1138 | 0.0224 |
| | Cer(d18:2/24:0) | -0.69124 | 0.0300 |
| | Cer(d18:2/26:1) | 0.75984 | 0.0003 |
| **Sphingomyelins** | SM(30:1) | 0.73017 | 0.0485 |
| | SM(32:1) | 0.85632 | 0.0143 |
| | SM(34:1) | 0.89942 | 0.0426 |
| | SM(38:1) | 1.1291 | 0.0296 |
| | SM(42:1) | -1.0103 | 0.0217 |
| | SM(42:4) | 1.0539 | 0.0424 |
| **Phospholipids** | | | |
| **Phosphatidic acids** | PA(34:2) | -0.57105 | 0.0130 |
| | PA(38:4) | 0.75981 | 0.0031 |
| **Phosphatidylcholines** | PC(34:1) | 1.0244 | 0.0078 |
| | PC(36:1) | 1.9891 | 0.0028 |
| | PC(36:3) | 0.89001 | 0.0171 |
| | PC(38:2) | 0.78019 | 0.0243 |
| | PC(38:3) | 1.9731 | 0.0071 |
| | PC(38:4) | 1.5209 | 0.0156 |
| | PC(38:5) | 1.1312 | 0.0088 |
| | PC(38:6) | 0.69253 | 0.0106 |
| | PC(40:4) | 0.99161 | 0.0198 |
| | PC(40:5) | 1.6062 | 0.0016 |
| | PC(40:6) | 1.3364 | 0.0083 |
| | PC(40:7) | 1.1116 | 0.0098 |
| **Phosphatidylethanolamines** | PE(32:1) | -1.1644 | 0.0455 |
| | PE(34:2) | -1.7663 | 0.0221 |
| | PE(36:2) | -0.44408 | 0.0409 |
| | PE(36:3) | -1.2293 | 0.0246 |
| | PE(36:5) | -1.1424 | 0.0208 |
| | PE(38:2) | -0.65029 | 0.0479 |
| | PE(38:5) | -0.6575 | 0.0495 |
| | PE(40:5) | -0.74282 | 0.0105 |
| | PE(40:7) | -0.7277 | 0.0405 |
| | PE(40:8) | -0.37791 | 0.0384 |
| **Phosphatidylglycerols** | PG(34:2) | -1.2053 | 0.0096 |
| | PG(38:3) | 1.1116 | 0.0400 |
| **Phosphatidylinositols** | PI(34:2) | -1.3178 | 0.0000 |
| | PI(36:1) | 0.61879 | 0.0059 |
| | PI(36:2) | -0.64978 | 0.0055 |
| | PI(38:2) | 2.1895 | 0.0030 |
| | PI(38:3) | 2.0998 | 0.0031 |
| | PI(38:4) | 0.80225 | 0.0163 |
| | PI(40:4) | 1.0617 | 0.0150 |
| | PI(40:5) | 0.66546 | 0.0223 |
| **Phosphatidylserines** | PS(38:4) | 0.63661 | 0.0219 |
| | PS(40:4) | 0.5581 | 0.0441 |
| **LysoPC** | LPC(16:0) | 0.63904 | 0.0181 |
| | LPC(16:1) | 0.65309 | 0.0260 |
| | LPC(18:0) | 1.1258 | 0.0093 |
| | LPC(18:1) | 1.5026 | 0.0111 |
| | LPC(20:0) | -1.4801 | 0.0157 |
| | LPC(20:1) | 0.52554 | 0.0372 |
| | LPC(20:3) | 1.958 | 0.0068 |
| | LPC(20:4) | 1.2664 | 0.0476 |
| | LPC(20:5) | 0.76202 | 0.0192 |
| | LPC(22:5) | 0.8625 | 0.0033 |
| | LPC(22:6) | 0.81978 | 0.0466 |
| **LysoPE** | LPE(18:2) | -0.83845 | 0.0214 |
| | LPE(18:3) | -0.7949 | 0.0050 |
| | LPE(20:5) | -0.44755 | 0.0238 |
| | LPE(22:1) | -0.6372 | 0.0429 |
| **PE Plasmalogens** | PE(16:0p/20:3) | 0.64188 | 0.0185 |
| | PE(16:0p/20:4) | 0.30424 | 0.0294 |
| | PE(18:0p/18:1) | 0.50645 | 0.0063 |
| | PE(18:0p/20:4) | 0.54511 | 0.0385 |

#### Table S4 : Detailed lipid species significantly enriched in AdEV compared to source VAT corresponding to Venn Diagram presented in Figure 4G

| Sample type | Lean sEV | Obese sEV | Obese lEV | Lean lEV | Lean lEV + Obese sEV | Lean sEV + Obese sEV | Lean lEV + Lean sEV | Obese lEV + Obese sEV | Lean lEV + Obese lEV + Obese sEV | Lean lEV + Lean sEV + Obese lEV | Lean sEV + Obese lEV + Obese sEV | Lean lEV + Lean sEV + Obese lEV + Obese sEV |
|---|---|---|---|---|---|---|---|---|---|---|---|---|
| **Number of lipid species retrieved** | **10** | **11** | **8** | **7** | **1** | **6** | **4** | **4** | **2** | **1** | **5** | **16** |
| **Lipid species enriched in AdEV subtypes** | Cer(d18:2/22:0) | PA(40:4) | Cer(d18:0/23:0) | Cer(d18:2/14:0) | Cer(d18:2/26:0) | Cer(d18:1/23:0) | Cer(d18:1/18:1) | LPC(20:1) | Cer(d18:2/23:1) | LPC(22:4) | Cer(d18:0/20:0) | Cer(d18:0/16:0) |
| | Cer(d18:2/23:0) | PC(34:0) | LPC(18:0) | Cer(d18:2/18:1) | | Cer(d18:1/24:0) | Cer(d18:1/19:0) | PC(30:2) | PG(34:3) | | Cer(d18:0/24:1) | Cer(d18:0/18:0) |
| | Cer(d18:2/24:0) | PC(36:2) | LPC(18:1) | LPE(18:0) | | Cer(d18:1/24:1) | Cer(d18:2/19:0) | SM(34:1) | | | Cer(d18:1/20:0) | Cer(d18:0/22:0) |
| | Cer(d18:2/24:1) | PE(32:0) | LPC(20:2) | PC(30:1) | | Cer(d18:2/20:0) | Cer(d18:2/24:2) | SM(38:2) | | | Cer(d18:1/22:0) | Cer(d18:1/14:0) |
| | LPE(20:4) | PG(36:1) | LPC(22:3) | PG(32:0) | | Cer(d18:2/21:0) | | | | | PG(36:3) | Cer(d18:1/16:0) |
| | LPE(22:4) | PG(38:2) | LPE(22:3) | PS(32:0) | | PG(38:3) | | | | | | Cer(d18:1/18:0) |
| | LPE(22:5) | SM(36:1) | PE(18:0p/18:3) | PS(36:5) | | | | | | | | Cer(d18:1/25:0) |
| | PA(32:0) | SM(36:2) | SM(42:3) | | | | | | | | | Cer(d18:1/26:0) |
| | PE(16:0p/22:4) | SM(38:1) | | | | | | | | | | Cer(d18:1/26:1) |
| | PS(34:1) | SM(40:1) | | | | | | | | | | Cer(d18:2/16:0) |
| | | SM(42:2) | | | | | | | | | | Cer(d18:2/18:0) |
| | | | | | | | | | | | | PC(28:0) |
| | | | | | | | | | | | | PC(30:0) |
| | | | | | | | | | | | | PC(32:0) |
| | | | | | | | | | | | | PC(34:1) |
| | | | | | | | | | | | | PG(36:4) |

**Table S5 : Detailed lipid species significantly depleted in AdEV compared to source VAT corresponding to Venn Diagram presented in Figure 4H**

| Sample type | Lean sEV | Obese sEV | Obese lEV | Lean lEV | Lean lEV + Obese sEV | Lean sEV + Obese sEV | Lean lEV + Lean sEV | Obese lEV + Obese sEV | Lean lEV + Obese lEV + Obese sEV | Lean lEV + Lean sEV + Obese lEV | Lean sEV + Obese lEV + Obese sEV | Lean lEV + Lean sEV + Obese lEV + Obese sEV |
|---|---|---|---|---|---|---|---|---|---|---|---|---|
| Number of lipid species retrieved | 4 | 5 | 8 | 4 | 1 | 6 | 5 | 6 | 4 | 1 | 2 | 17 |
| Lipid species depleted in AdEV subtypes | LPC(16:0)<br>PC(34:3)<br>SM(32:1)<br>SM(38:1) | LPC(20:3)<br>LPC(22:6)<br>PE(18:0p/18:2)<br>PE(18:0p/20:5)<br>PI(36:1) | PA(38:5)<br>PC(36:0)<br>PC(36:6)<br>PE(16:0p/22:6)<br>PE(40:5)<br>PI(34:1)<br>PS(36:3)<br>PS(38:2) | PC(38:2)<br>PE(38:2)<br>SM(40:1)<br>SM(42:4) | PI(36:3) | PE(34:1)<br>PE(34:2)<br>SM(38:2)<br>SM(40:2)<br>SM(41:1)<br>SM(41:2) | PC(38:4)<br>PC(40:4)<br>PC(40:8)<br>PE(18:0p/22:6)<br>PE(34:0) | PE(28:0)<br>PE(38:3)<br>PE(40:8)<br>PI(38:3)<br>PI(38:6)<br>PI(40:6) | PE(36:2)<br>PI(36:4)<br>PI(38:4)<br>PI(38:5) | PC(40:7) | PE(32:1)<br>PI(36:2) | PC(36:3)<br>PC(36:4)<br>PC(36:5)<br>PC(38:3)<br>PC(38:5)<br>PC(38:6)<br>PC(40:5)<br>PC(40:6)<br>PE(36:3)<br>PE(36:4)<br>PE(36:5)<br>PE(38:4)<br>PE(38:5)<br>PE(38:6)<br>PE(40:6)<br>PE(40:7)<br>PS(40:6) |

**Table S3 : Plasma lipid fingerprints of lean and obese mice**

| | | obese *vs* lean | |
|---|---|---|---|
| | | Log2FC | *p* value |
| **Neutral Lipids** | | | |
| **Cholesterol esters** | CE(16:1) | 0.92818 | 0.0036 |
| | CE(18:1) | 1.106 | 0.0245 |
| | CE(18:3) | 0.27607 | 0.0088 |
| | CE(20:3) | 1.6667 | 0.0071 |
| | CE(20:4) | 1.4249 | 0.0088 |
| | CE(22:5) | 1.6288 | 0.0058 |
| | CE(22:6) | 1.1984 | 0.0054 |
| **Diacylglycerols** | DG(16:0_18:2) | -1.7555 | 0.0118 |
| | DG(16:1_18:1) | -1.2459 | 0.0163 |
| | DG(16:1_18:2) | -3.2715 | 0.0012 |
| | DG(18:2_18:2) | -2.5228 | 0.0049 |
| | DG(18:1_20:3) | 1.2761 | 0.0142 |
| | DG(16:0_22:6) | -1.0062 | 0.0492 |
| | DG(18:2_20:4) | -2.5898 | 0.0116 |
| | DG(18:1_22:6) | -1.8364 | 0.0121 |
| **Triacylglycerols** | TG(48:0-16:0_32:0) | -1.1351 | 0.0488 |
| | TG(48:1-16:1_32:0) | -1.4872 | 0.0362 |
| | TG(48:2-14:1_34:1) | -1.1552 | 0.0379 |
| | TG(48:2-16:0_32:2) | -2.2711 | 0.0033 |
| | TG(48:2-18:1_30:1) | -1.4775 | 0.0154 |
| | TG(48:2-18:2_30:0) | -2.9261 | 0.0030 |
| | TG(48:3-14:0_34:3) | -3.5451 | 0.0045 |
| | TG(48:3-16:1_32:2) | -2.4038 | 0.0043 |
| | TG(49:1-14:0_35:1) | -1.1129 | 0.0040 |
| | TG(49:1-15:0_34:1) | -1.7034 | 0.0140 |
| | TG(50:2-18:0_32:2) | -2.1336 | 0.0025 |
| | TG(50:2-18:1_32:1) | -1.1721 | 0.0143 |
| | TG(50:2-18:2_32:0) | -2.6086 | 0.0040 |
| | TG(50:3-18:1_32:2) | -1.8793 | 0.0132 |
| | TG(50:3-18:2_32:1) | -3.599 | 0.0006 |
| | TG(50:4-14:0_36:4) | -3.5617 | 0.0015 |
| | TG(50:4-18:2_32:2) | -3.522 | 0.0016 |
| | TG(51:1-18:1_33:0) | -1.4596 | 0.0056 |
| | TG(51:2-15:0_36:2) | -1.9691 | 0.0183 |
| | TG(51:2-16:0_35:2) | -1.8693 | 0.0049 |
| | TG(52:3-18:2_34:1) | -2.7666 | 0.0009 |
| | TG(52:4-16:0_36:4) | -3.4432 | 0.0008 |
| | TG(52:4-16:1_36:3) | -2.9304 | 0.0057 |
| | TG(52:4-20:4_32:0) | -0.99352 | 0.0347 |
| | TG(52:5-20:4_32:1) | -2.0387 | 0.0064 |
| | TG(54:1-18:1_36:0) | -2.2306 | 0.0344 |
| | TG(54:4-18:0_36:4) | -2.6158 | 0.0043 |
| | TG(54:4-18:2_36:2) | -2.2552 | 0.0042 |
| | TG(54:5-18:1_36:4) | -2.839 | 0.0020 |
| | TG(54:6-18:2_36:4) | -2.8723 | 0.0065 |
| | TG(54:6-18:3_36:3) | -3.0976 | 0.0024 |
| | TG(54:6-20:4_34:2) | -1.9806 | 0.0038 |
| | TG(54:6-22:6_32:0) | -2.2953 | 0.0478 |
| | TG(56:6-20:5_36:1) | -1.9816 | 0.0018 |
| | TG(56:6-22:5_34:1) | -1.4341 | 0.0105 |
| | TG(56:8-20:4_36:4) | -2.1677 | 0.0021 |
| | TG(58:8-22:6_36:2) | -1.7758 | 0.0159 |
| **Sphingolipids** | | | |
| **Ceramides** | Cer(d18:0/18:0) | 0.86002 | 0.0499 |
| | Cer(d18:0/18:1) | 0.83728 | 0.0468 |
| | Cer(d18:0/20:0) | 0.94519 | 0.0342 |
| | Cer(d18:0/24:1) | 1.4545 | 0.0284 |
| | Cer(d18:1/14:0) | 2.48 | 0.0217 |
| | Cer(d18:1/16:0) | 1.739 | 0.0154 |
| | Cer(d18:1/18:0) | 4.2423 | 0.0273 |
| | Cer(d18:1/18:1) | 2.0894 | 0.0198 |
| | Cer(d18:1/19:0) | 1.3885 | 0.0223 |
| | Cer(d18:1/20:0) | 2.2942 | 0.0007 |
| | Cer(d18:1/23:0) | 0.45342 | 0.0434 |
| | Cer(d18:1/24:1) | 1.475 | 0.0060 |

| | | | |
|---|---|---|---|
| | Cer(d18:1/26:0) | 1.1633 | 0.0211 |
| | Cer(d18:1/26:1) | 1.1369 | 0.0148 |
| | Cer(d18:2/16:0) | 0.96748 | 0.0121 |
| | Cer(d18:2/18:0) | 2.5047 | 0.0135 |
| | Cer(d18:2/18:1) | 1.0745 | 0.0097 |
| | Cer(d18:2/20:1) | 1.1138 | 0.0224 |
| | Cer(d18:2/24:0) | -0.69124 | 0.0300 |
| | Cer(d18:2/26:1) | 0.75984 | 0.0003 |
| **Sphingomyelins** | SM(30:1) | 0.73017 | 0.0485 |
| | SM(32:1) | 0.85632 | 0.0143 |
| | SM(34:1) | 0.89942 | 0.0426 |
| | SM(38:1) | 1.1291 | 0.0296 |
| | SM(42:1) | -1.0103 | 0.0217 |
| | SM(42:4) | 1.0539 | 0.0424 |
| **Phospholipids** | | | |
| **Phosphatidic acids** | PA(34:2) | -0.57105 | 0.0130 |
| | PA(38:4) | 0.75981 | 0.0031 |
| **Phosphatidylcholines** | PC(34:1) | 1.0244 | 0.0078 |
| | PC(36:1) | 1.9891 | 0.0028 |
| | PC(36:3) | 0.89001 | 0.0171 |
| | PC(38:2) | 0.78019 | 0.0243 |
| | PC(38:3) | 1.9731 | 0.0071 |
| | PC(38:4) | 1.5209 | 0.0156 |
| | PC(38:5) | 1.1312 | 0.0088 |
| | PC(38:6) | 0.69253 | 0.0106 |
| | PC(40:4) | 0.99161 | 0.0198 |
| | PC(40:5) | 1.6062 | 0.0016 |
| | PC(40:6) | 1.3364 | 0.0083 |
| | PC(40:7) | 1.1116 | 0.0098 |
| **Phosphatidylethanolamines** | PE(32:1) | -1.1644 | 0.0455 |
| | PE(34:2) | -1.7663 | 0.0221 |
| | PE(36:2) | -0.44408 | 0.0409 |
| | PE(36:3) | -1.2293 | 0.0246 |
| | PE(36:5) | -1.1424 | 0.0208 |
| | PE(38:2) | -0.65029 | 0.0479 |
| | PE(38:5) | -0.6575 | 0.0495 |
| | PE(40:5) | -0.74282 | 0.0105 |
| | PE(40:7) | -0.7277 | 0.0405 |
| | PE(40:8) | -0.37791 | 0.0384 |
| **Phosphatidylglycerols** | PG(34:2) | -1.2053 | 0.0096 |
| | PG(38:3) | 1.1116 | 0.0400 |
| **Phosphatidylinositols** | PI(34:2) | -1.3178 | 0.0000 |
| | PI(36:1) | 0.61879 | 0.0059 |
| | PI(36:2) | -0.64978 | 0.0055 |
| | PI(38:2) | 2.1895 | 0.0030 |
| | PI(38:3) | 2.0998 | 0.0031 |
| | PI(38:4) | 0.80225 | 0.0163 |
| | PI(40:4) | 1.0617 | 0.0150 |
| | PI(40:5) | 0.66546 | 0.0223 |
| **Phosphatidylserines** | PS(38:4) | 0.63661 | 0.0219 |
| | PS(40:4) | 0.5581 | 0.0441 |
| **LysoPC** | LPC(16:0) | 0.63904 | 0.0181 |
| | LPC(16:1) | 0.65309 | 0.0260 |
| | LPC(18:0) | 1.1258 | 0.0093 |
| | LPC(18:1) | 1.5026 | 0.0111 |
| | LPC(20:0) | -1.4801 | 0.0157 |
| | LPC(20:1) | 0.52554 | 0.0372 |
| | LPC(20:3) | 1.958 | 0.0068 |
| | LPC(20:4) | 1.2664 | 0.0476 |
| | LPC(20:5) | 0.76202 | 0.0192 |
| | LPC(22:5) | 0.8625 | 0.0033 |
| | LPC(22:6) | 0.81978 | 0.0466 |
| **LysoPE** | LPE(18:2) | -0.83845 | 0.0214 |
| | LPE(18:3) | -0.7949 | 0.0050 |
| | LPE(20:5) | -0.44755 | 0.0238 |
| | LPE(22:1) | -0.6372 | 0.0429 |
| **PE Plasmalogens** | PE(16:0p/20:3) | 0.64188 | 0.0185 |
| | PE(16:0p/20:4) | 0.30424 | 0.0294 |
| | PE(18:0p/18:1) | 0.50645 | 0.0063 |
| | PE(18:0p/20:4) | 0.54511 | 0.0385 |

#### Table S4 : Detailed lipid species significantly enriched in AdEV compared to source VAT corresponding to Venn Diagram presented in Figure 4G

| **Sample type** | **Lean sEV** | **Obese sEV** | **Obese lEV** | **Lean lEV** | **Lean lEV + Obese sEV** | **Lean sEV + Obese sEV** | **Lean lEV + Lean sEV** | **Obese lEV + Obese sEV** | **Lean lEV + Obese lEV + Obese sEV** | **Lean lEV + Lean sEV + Obese lEV** | **Lean sEV + Obese lEV + Obese sEV** | **Lean lEV + Lean sEV + Obese lEV + Obese sEV** |
|---|---|---|---|---|---|---|---|---|---|---|---|---|
| **Number of lipid species retrieved** | **10** | **11** | **8** | **7** | **1** | **6** | **4** | **4** | **2** | **1** | **5** | **16** |
| **Lipid species enriched in AdEV subtypes** | Cer(d18:2/22:0)<br>Cer(d18:2/23:0)<br>Cer(d18:2/24:0)<br>Cer(d18:2/24:1)<br>LPE(20:4)<br>LPE(22:4)<br>LPE(22:5)<br>PA(32:0)<br>PE(16:0p/22:4)<br>PS(34:1) | PA(40:4)<br>PC(34:0)<br>PC(36:2)<br>PE(32:0)<br>PG(36:1)<br>PG(38:2)<br>SM(36:1)<br>SM(36:2)<br>SM(38:1)<br>SM(40:1)<br>SM(42:2) | Cer(d18:0/23:0)<br>LPC(18:0)<br>LPC(18:1)<br>LPC(20:2)<br>LPC(22:3)<br>LPE(22:3)<br>PE(18:0p/18:3)<br>SM(42:3) | Cer(d18:2/14:0)<br>Cer(d18:2/18:1)<br>LPE(18:0)<br>PC(30:1)<br>PG(32:0)<br>PS(32:0)<br>PS(36:5) | Cer(d18:2/26:0) | Cer(d18:1/23:0)<br>Cer(d18:1/24:0)<br>Cer(d18:1/24:1)<br>Cer(d18:2/20:0)<br>Cer(d18:2/21:0)<br>PG(38:3) | Cer(d18:1/18:1)<br>Cer(d18:1/19:0)<br>Cer(d18:2/19:0)<br>Cer(d18:2/24:2) | LPC(20:1)<br>PC(30:2)<br>SM(34:1)<br>SM(38:2) | Cer(d18:2/23:1)<br>PG(34:3) | LPC(22:4) | Cer(d18:0/20:0)<br>Cer(d18:0/24:1)<br>Cer(d18:1/20:0)<br>Cer(d18:1/22:0)<br>PG(36:3) | Cer(d18:0/16:0)<br>Cer(d18:0/18:0)<br>Cer(d18:0/22:0)<br>Cer(d18:1/14:0)<br>Cer(d18:1/16:0)<br>Cer(d18:1/18:0)<br>Cer(d18:1/25:0)<br>Cer(d18:1/26:0)<br>Cer(d18:1/26:1)<br>Cer(d18:2/16:0)<br>Cer(d18:2/18:0)<br>PC(28:0)<br>PC(30:0)<br>PC(32:0)<br>PC(34:1)<br>PG(36:4) |

**Table S5 : Detailed lipid species significantly depleted in AdEV compared to source VAT corresponding to Venn Diagram presented in Figure 4H**

| Sample type | Lean sEV | Obese sEV | Obese lEV | Lean lEV | Lean lEV + Obese sEV | Lean sEV + Obese sEV | Lean lEV + Lean sEV | Obese lEV + Obese sEV | Lean lEV + Obese lEV + Obese sEV | Lean lEV + Lean sEV + Obese lEV | Lean sEV + Obese lEV + Obese sEV | Lean lEV + Lean sEV + Obese lEV + Obese sEV |
|---|---|---|---|---|---|---|---|---|---|---|---|---|
| Number of lipid species retrieved | 4 | 5 | 8 | 4 | 1 | 6 | 5 | 6 | 4 | 1 | 2 | 17 |
| Lipid species depleted in AdEV subtypes | LPC(16:0)<br>PC(34:3)<br>SM(32:1)<br>SM(38:1) | LPC(20:3)<br>LPC(22:6)<br>PE(18:0p/18:2)<br>PE(18:0p/20:5)<br>PI(36:1) | PA(38:5)<br>PC(36:0)<br>PC(36:6)<br>PE(16:0p/22:6)<br>PE(40:5)<br>PI(34:1)<br>PS(36:3)<br>PS(38:2) | PC(38:2)<br>PE(38:2)<br>SM(40:1)<br>SM(42:4) | PI(36:3) | PE(34:1)<br>PE(34:2)<br>SM(38:2)<br>SM(40:2)<br>SM(41:1)<br>SM(41:2) | PC(38:4)<br>PC(40:4)<br>PC(40:8)<br>PE(18:0p/22:6)<br>PE(34:0) | PE(28:0)<br>PE(38:3)<br>PE(40:8)<br>PI(38:3)<br>PI(38:6)<br>PI(40:6) | PE(36:2)<br>PI(36:4)<br>PI(38:4)<br>PI(38:5) | PC(40:7) | PE(32:1)<br>PI(36:2) | PC(36:3)<br>PC(36:4)<br>PC(36:5)<br>PC(38:3)<br>PC(38:5)<br>PC(38:6)<br>PC(40:5)<br>PC(40:6)<br>PE(36:3)<br>PE(36:4)<br>PE(36:5)<br>PE(38:4)<br>PE(38:5)<br>PE(38:6)<br>PE(40:6)<br>PE(40:7)<br>PS(40:6) |

#### KEY RESOURCES TABLE

| REAGENT or RESOURCE | SOURCE | IDENTIFIER |
|---|---|---|
| ***Antibodies*** | | |
| CD9 | BD Biosciences | Cat #553758 |
| CD63 | MBL international | Cat #D263-3 |
| Flotillin-2 | BD Biosciences | Cat #9101S |
| Grp94/Endoplasmin | Enzo life Sciences | Cat #ADI-SPA-850 |
| Perilipin-1 | Progen | Cat #GP29 |
| IRDye® 800CW and 680RD secondary antibody (anti-mouse and anti-rabbit) | LI-COR Biosciences | Cat #926-32211, #926-32210, #926-68071, #926-68070 |
| ***Chemicals, peptides, and recombinant proteins*** | | |
| All internal lipid standards for quantitative lipidomics | Avanti® Polar Lipids | https://avantilipids.com/divisions/lipidomics/lipid-maps-ms-standards |
| DMEM 4.5g/L Glucose | Gibco part of Thermofisher Scintific | Cat #41966-029 |
| 4X Laemmli Sample Buffer | Bio-Rad | Cat #161-0747 |
| Odyssey blocking buffer (TBS) | LI-COR Biosciences | Cat #927-50010 |
| ***Deposited data*** | | |
| Lipidomic datasets of AdEV and tissue analyzed | This paper | Supplemental Tables |
| ***Experimental models: Organisms/strains*** | | |
| B6.Cg-Lepob/J | Charles River (JAX™ mice strain) | Stock number #000632 |
| ***Software and algorithms*** | | |
| Multi Experiment Viewer (MeV) software version 4.9 | | https://sourceforge.net/projects/mev-tm4/ |
| MetaboAnalyst open source software | | |
| GraphPad Prism | GraphPad Software, Inc | https://www.graphpad.com/ |
| Image Studio software | LI-COR Biosciences | https://www.licor.com/bio/products/software/image_studio/ |
| Image J | Downloaded from https://imagej.nih.gov/ij/ | https://imagej.nih.gov/ij/ |

| | | |
|---|---|---|
| Nanosight NTA software (version 3.1) | Malvern Panalytical | https://www.malvernpanalytical.com/en/support/product-support/software/NanoSight-NTA-software-update-v3-10-l2 |
